# scNODE : Generative Model for Temporal Single Cell Transcriptomic Data Prediction

**DOI:** 10.1101/2023.11.22.568346

**Authors:** Jiaqi Zhang, Erica Larschan, Jeremy Bigness, Ritambhara Singh

## Abstract

Measurement of single-cell gene expression at different timepoints enables the study of cell development. However, due to the resource constraints and technical challenges associated with the single-cell experiments, researchers can only profile gene expression at discrete and sparsely-sampled timepoints. This missing timepoint information impedes downstream cell developmental analyses. We propose `scNODE`, an end-to-end deep learning model that can predict *in silico* single-cell gene expression at unobserved timepoints. `scNODE` integrates a variational autoencoder (VAE) with neural ordinary differential equations (ODEs) to predict gene expression using a continuous and non-linear latent space. Importantly, we incorporate a dynamic regularization term to learn a latent space that is robust against distribution shifts when predicting single-cell gene expression at unobserved timepoints. Our evaluations on three real-world scRNA-seq datasets show that `scNODE` achieves higher predictive performance than state-of-the-art methods. We further demonstrate that `scNODE`’s predictions help cell trajectory inference under the missing timepoint paradigm and the learned latent space is useful for *in silico* perturbation analysis of relevant genes along a developmental cell path. The data and code are publicly available at https://github.com/rsinghlab/scNODE.

## 1 Introduction

A fundamental challenge in biology is understanding how gene expression changes over time during cell development (Moris et al., 2016; Fleck et al., 2022). With the advent of high-throughput single-cell RNA sequencing (scRNA-seq), researchers now routinely profile gene expression at single-cell resolution (Hwang et al., 2018; Chen et al., 2019), revealing substantial heterogeneity within the same tissue type or developmental stage (Chen et al., 2022a; Ding et al., 2022). Profiling scRNA-seq data at multiple timepoints allows us to understand how biological processes unfold within cell populations by inferring cellular trajectories, identifying cell fate transitions, and characterizing differentially expressed genes (Qiu et al., 2022; Liu et al., 2022).

Temporal scRNA-seq experiments, however, have significant limitations that impede developmental studies. Due to the substantial time and resources involved in conducting these experiments, researchers generally only profile gene expression at discrete and sparsely-sampled timepoints (Ding et al., 2022). Since it is infeasible to make continuous-time observations, this results in information loss between consecutively measured discrete timepoints. Therefore, it is crucial to develop a model that predicts realistic *in silico* gene expression at any timepoint, whether between (interpolation) or beyond the measured time intervals (extrapolation), to enable single-cell temporal analyses (Fig. 1**A-B**).

**Figure 1:**
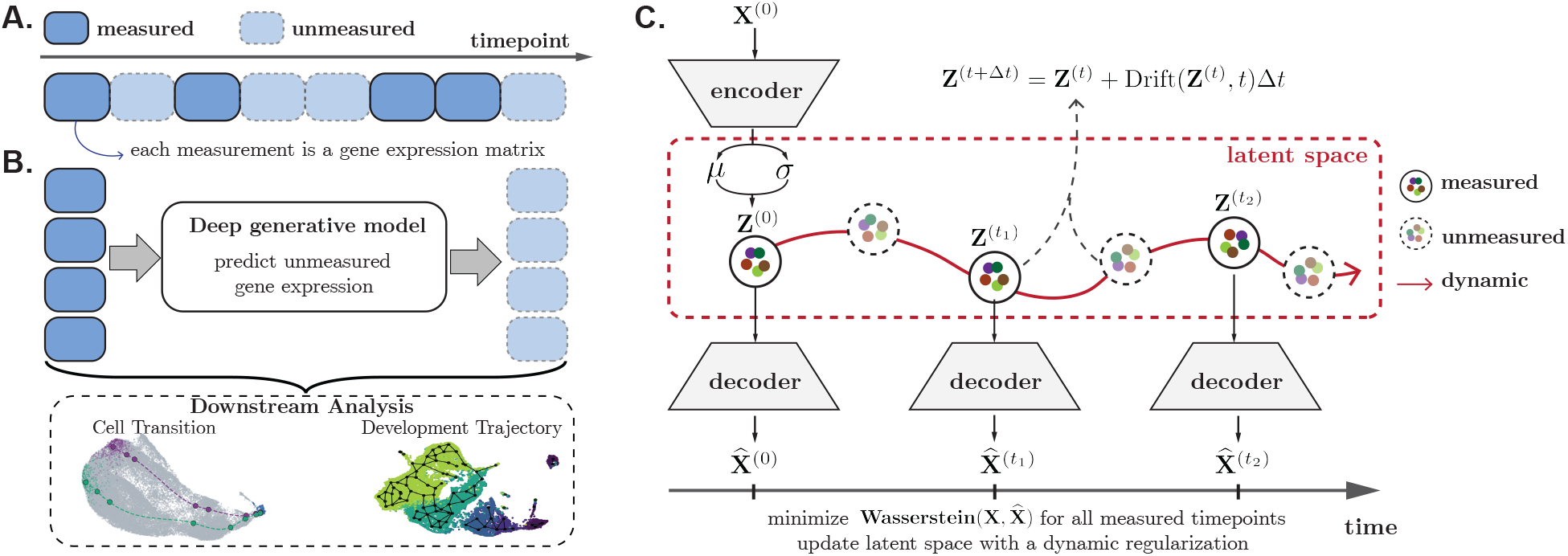
Model overview. (**A**) The measurements in temporally recorded single-cell data are sparse and unaligned, such that some timepoints are not measured and measurements at different timepoints contain different sets of cells. The data sets in this study sample cellular populations at uniformly spaced time points, although uniform spacing is not a requirement (as depicted in this graphic). (**B**) `scNODE` is used to predict expression at unmeasured timepoints. Predictions can be used to analyze cell state transitions and developmental trajectories. (**C**) `scNODE` uses VAEs to find an optimal low-dimensional representation. `scNODE` then uses this representation to solve neural ODEs and predict gene expression at any timepoint *t* = *t*_1_, *t*_2_, … starting from the first timepoint *t* = 0. Optimal models are fitted by minimizing the Wasserstein distance between observations and predictions at measured timepoints. `scNODE` also introduces a dynamic regularization to learn a latent space robust to distribution shift.

Single-cell gene expression prediction has been investigated in previous studies (Cao et al., 2021) through various generative models. Existing methods use generative adversarial networks (GANs) (Marouf et al., 2020) or statistical frameworks (Zappia et al., 2017; Risso et al., 2018; Li and Li, 2019; Sun et al., 2021) to fit characteristics of the observed single-cell gene expression and augment the dataset. However, these methods are designed for single-cell gene expression measured at a single time snapshot and do not consider developmental dynamics.

Several computational methods have been developed to learn gene expression dynamics by inferring cell trajectories (Trapnell et al., 2014; Qiu et al., 2017; Herman et al., 2018; Saelens et al., 2019; Wolf et al., 2019; Li, 2023) from temporal scRNA-seq. However, most of them infer pseudo-times, which analyze cell differentiation based on geometric distances in low-dimensional spaces. Hence, they do not accurately model how cells develop in real physical time. Sagittarius method (Woicik et al., 2023) predicts gene expressions at future timepoints but requires cell matching between timepoints. This matching is hard to obtain for real-world scRNA-seq as the cells are lysed during the experiment. RNA velocity based methods (La Manno et al., 2018; Bergen et al., 2021, 2020) are used to uncover developmental trends and underlying kinetics of gene expression. These methods require deeply sequenced datasets to obtain spliced and unspliced counts as prior information, and most of them are non-generative methods. Waddington-OT (Schiebinger et al., 2019) and TrajectoryNet (Tong et al., 2020) interpolate single-cell gene expression between two timepoints in a developmental trajectory but are unable to extrapolate beyond the observed timepoints.

Recently, a few generative methods have modeled cell differentiation trajectory and predicted gene expression at multiple unobserved time points. For example, PRESCIENT (Yeo et al., 2021) applies principal component analysis (PCA) to reduce high-dimensional gene space to a low-dimensional representation. It then applies neural ordinary differential equations (ODEs) (Chen et al., 2018) in this low-dimensional space to model the cell developmental trajectory. Multiple studies (Ding et al., 2018; Xiang et al., 2021; Tran et al., 2021) have found that PCA-based low-dimensional representations of scRNA-seq expression cannot capture complex relationships in the highly heterogeneous single-cell data. Thus, PCA may have the issue of overcrowding representation (Tran et al., 2021; Kobak and Berens, 2019), where cells of different types are poorly separated in PCA-based low-dimensional space, and cellular variations are lost. MIOFlow (Huguet et al., 2022) replaces PCA with a geodesic variational autoencoder (VAE) to learn non-linear low-dimensional representations that better retain cellular variations. However, this method fixes its low-dimensional space at the beginning, followed by neural ODE modeling. We hypothesize that a fixed low-dimensional representation obtained using observed timepoints may not generalize to an unobserved timepoint, if its underlying distribution differs substantially.

We propose a single-cell neural ODE (`scNODE`) framework, which integrates a VAE (Kingma et al., 2019) with neural ODE to predict accurate gene expression at any unmeasured timepoint using a dynamic low-dimensional space (Fig. 1**C**). First, `scNODE` uses VAE to obtain a low-dimensional representation that retains cellular variations. `scNODE` then predicts single-cell gene expression for unmeasured timepoints with a neural ODE. Importantly, `scNODE` uses a *dynamic regularization* term to incorporate the dynamic manifold learned from neural ODE as a prior. Therefore, the learned VAE space is robust to the distribution shift between observed and unobserved time points. We evaluate `scNODE` on three real-world scRNA-seq datasets. Our results demonstrate that `scNODE` predicts single-cell gene expression more accurately than state-of-the-art methods for unobserved timepoints (interpolation and extrapolation). We further show that `scNODE` gene expression predictions assist with cell developmental trajectory inference. Finally, `scNODE` learns an interpretable low-dimensional space, which enables conducting *in silico* perturbation analysis of relevant genes to study cell development.

## 2 Method

*Notation:* 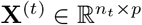 denote the gene expression matrix at the *t*-th time point of *n*_*t*_ cells by *p* genes. Given gene expression measurements at observed timepoints *{***X**^(*t*)^*}*_*t*∈*𝒯*_, our goal is to predict gene expression at any timepoint. Here, *𝒯* ⊆ {0, 1, 2, … } denotes the observed timepoint indices. Note that we assume the same set of highly variable genes (HVGs) is measured at each time point, but due to the destruction of cells during scRNA experiments, a different set of cells is measured at each time point. Furthermore, let 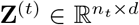 denote the *d*-dimensional (*d* ≪ *p*) latent variable at the *t*-th time point.

### 2.1 scNODE uses VAE to learn complex low-dimensional space

`scNODE` uses a Variational Autoencoder (VAE) to learn a low-dimensional representation (also known as latent space) of the single-cell gene expression measurements at the observed timepoints. A VAE is a neural network-based generative model that maps high-dimensional data to a low-dimensional representation and is widely used in many single-cell studies (Grønbech et al., 2020; Svensson et al., 2020; Wang and Gu, 2018; Lin et al., 2020), showing superior performance. The benefit of using VAEs for single-cell data over PCA is that the non-linearity of neural networks can more effectively capture complex cell relationships and cellular variations.

`scNODE` first pre-trains a VAE to obtain a latent space that captures the gene expression of single cells at all available training timepoints. Specifically, `scNODE` trains the VAE component to perform multi-timepoint modeling by inputting the gene expression of all observed cells **X**_ALL_ = CONCAT ( **X**^(*t*)^ | *t* ∈ *𝒯*) . VAE is composed of two neural networks: (1) the encoder network Enc_*ϕ*_ maps expression from gene space ℝ^*p*^ to a low-dimensional latent space ℝ^*d*^, parameterized by a Gaussian distribution *𝒩* (*µ, σ*), and (2) the decoder network Dec_*θ*_ maps samples in the latent space to gene space in the opposite direction to reconstruct the input. Given **X**_ALL_, VAE learns latent representations through

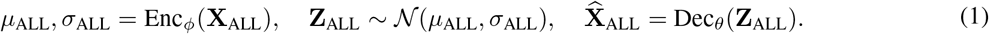

The encoder and decoder networks are parameterized by *ϕ* and *θ*, correspondingly. VAE minimizes a combination of the (1) mean squared error (MSE) between input gene expression and the reconstructed gene expression from the decoder and (2) the Kulback-Leibler (KL) divergence (Csiszár, 1975) between the latent distribution and a standard Gaussian prior:

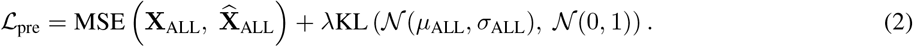

KL divergence is an information-theoretic quantity that quantifies the distance between two probability distributions. By using a Gaussian prior, the KL divergence forces the encoder to find latent representations of the cells that are well-separated. `scNODE` pre-trains the VAE component with Eq. 2, such that the encoder latent space preserves the variation of all observed cells. Next, we model the developmental dynamics of this latent space.

### 2.2 scNODE uses neural ODE to model cell dynamics

`scNODE` uses ordinary differential equations (ODEs) to model cell developmental dynamics in the latent space learned by the VAE. An ODE is an equation describing how a quantity *x* changes with respect to an independent variable *y*, such that d*x* = *f* (*x*; *y*)d*y* where function *f* represents the derivative. Therefore, we can use differential equations to model how gene expression changes with respect to time. However, for the high-dimensional data, finding the solution of the derivative function *f* through numerical methods is intractable and computationally expensive (Kidger, 2022). Therefore, recent studies adopt neural networks to approximate the derivative function and have proposed neural ODEs (Chen et al., 2018). Neural ODEs (formulated with respect to time) are useful for constructing continuous-time trajectories and have been explored before to model single-cell development (Matsumoto et al., 2017; Qiu et al., 2022; Chen et al., 2022b).

`scNODE` uses neural ODEs to parameterize the continuous dynamics of gene expression in the latent space. Specifically, `scNODE` quantifies changes of cell latent representation **Z**^(*t*)^ (from VAE encoding) at time *t* through a neural ODE

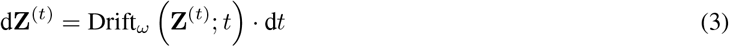

Here, Drift_*ω*_ is a non-linear neural network with parameterization *ω*, modeling the developmental velocities in the latent space, such that Drift_*ω*_(**Z**^(*t*)^; *t*) represents the direction and strength of cellular transitions. `scNODE` computes the latent representation **Z**^(0)^ of cells at the first timepoint *t* = 0 through the encoder network (pre-trained in the previous step) and predicts the subsequent cell states step-wise at any timepoint *t* as follows:

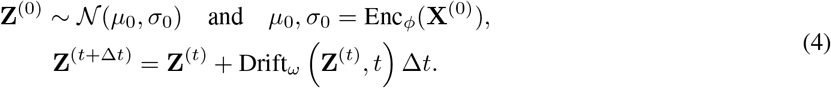

Here, hyperparameter Δ*t* denotes step size and drift term Drift_*ω*_ **Z**^(*t*)^, *t* Δ*t* represents the forward steps taken in the latent space. While there are several methods that can solve this ODE, in this work, we apply the commonly used first-order Euler method (in Eq. 4) for convenience of explanation. However, in our implementation, one can specify any ODE solver.

To fit the continuous trajectory (controlled by Drift_*ω*_) to the observations, `scNODE` minimizes the difference between the input and the reconstructed gene expression. Specifically, at each measured timepoint *t* ∈ *𝒯*, `scNODE` uses the decoder Dec_*θ*_ to convert latent variables **Z**^(*t*)^ generated from Eq. 4 back to the gene space through 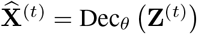. Because we have no correspondence between true cells and cells generated from the ODE model, `scNODE` utilizes the Wasserstein metric (Cuturi, 2013) to measure the distance between distributions defined by ground truth **X** and predictions 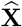 as

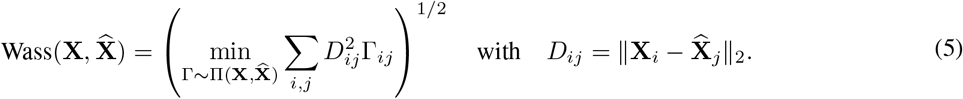

Here, 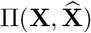 denotes the set of all transport plans between each cell of **X** and 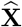 and *D*_*ij*_ represents the *𝓁*_2_ distance, such that the Wasserstein metric adopts the minimal-cost transport plan Γ to measure the data dissimilarity.

### 2.3 scNODE uses dynamic regularization for learning a robust latent space

The VAE and its corresponding latent space are trained using training (or observed) timepoints. In the temporal scRNA-seq data where gene expression distribution changes substantially, a fixed low-dimensional representation obtained using training timepoints may not generalize to a testing (or unobserved) timepoint. To overcome this distribution shift issue, `scNODE` uses a **dynamic regularization** term to update the VAE space dynamically such that it captures both cellular variations and the developmental dynamics of the scRNA-seq data. Specifically, `scNODE` minimizes the difference between the latent variables generated by the VAE (i.e., Enc_*ϕ*_(**X**)) and the dynamics learned by the ODE (i.e., **Z**). Because we have no correspondence between them, `scNODE` again uses Wasserstein distance to evaluate their difference at each observed timepoint *t* ∈ *𝒯* and defines the dynamic regularization as

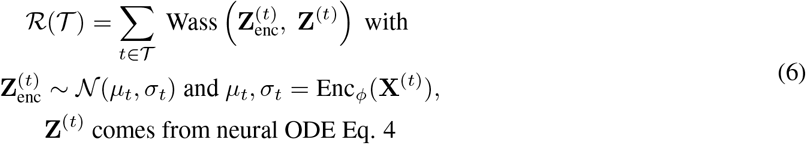

Therefore, `scNODE` jointly updates VAE and neural ODE, optimizing parameters *ϕ, θ*, and *ω*, by minimizing the regularized loss function

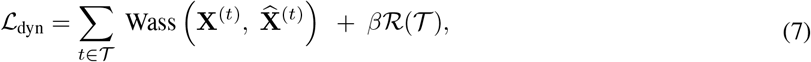

so that the overall dynamics update the final latent space of `scNODE` through dynamic regularization and corresponding hyperparameter *β*.

Our dynamic regularization improves upon previous generative models (Schiebinger et al., 2019; Tong et al., 2020; Huguet et al., 2022; Yeo et al., 2021), which fix the latent space at the beginning and then learn the cell dynamics. `scNODE` uses the dynamic manifold learned from neural ODE as a prior and enforces VAE latent space to incorporate it using dynamic regularization. Therefore, updating the VAE ensures that `scNODE` fits the data better and learns a latent space that is robust to distribution shift when generating gene expression for unobserved timepoints. Previous work by Connor et al. (2021) similarly introduced a linear dynamic system-based model to learn a smooth low-dimensional manifold and then incorporated this manifold (as a prior) into VAE to encourage the latent space to fit the data manifold better. This resolved the issue that VAE latent representations sometimes would not match the true data manifold and poorly defined natural paths between data points. Therefore, our dynamic regularization will result in improved representation learning.

## 3 Results

### 3.1 Experimental Setup

#### Datasets

We use three publicly available scRNA-seq datasets to demonstrate the capabilities of `scNODE` in predicting developmental dynamics from real-world single-cell gene expression data. These datasets are summarized in Table 1, have more than 10 timepoints, and cover various species and tissues. For each timepoint, gene expression is measured at a developmental stage (ZB), every hour (DR), or every 12 hours (SC). To make computations tractable, we relabel timepoints with consecutive natural numbers starting from 0. In particular, the meaning of interpolating or extrapolating is defined by these data-specific units. For example, extrapolating one timepoint in the DR dataset means predicting gene expressions for the next hour. We use the data after removing batch effects among different timepoints. In each experiment, we select the top 2000 most highly variable genes (HVGs) from the datasets and normalize the unique molecular identifier (UMI) count expression through a log transformation with pseudo-count.

**Table 1:** Data descriptions of the three real-world scRNA-seq datasets used in experiments.

| ID | Dataset | Species | # Cells | # Timepoints | Source |
| --- | --- | --- | --- | --- | --- |
| ZB | zebrafish embryo | *Danio rerio* | 38731 | 12 | Broad Single-Cell Portal SCP162 (Farrell et al., 2018) |
| DR | drosophila | *Drosophila melanogaster* | 27386 | 11 | GSE190149 (Calderon et al., 2022) |
| SC | Schiebinger2019 | *Mus musculus* | 236285 | 19 | (Schiebinger et al., 2019) |

#### Training and testing

We test `scNODE` ‘s performance in predicting gene expression at unobserved timepoints. Specifically, for each dataset, we remove several timepoints to test whether `scNODE` can recover these left-out observations. We design three tasks: (1) **easy** tasks where we remove uniformly spaced timepoints in the middle of the measured time range for interpolating, (2) **medium** task where we remove the last few timepoints for extrapolating beyond the measured time range, and (3) **hard** tasks where we combine the interpolation and extrapolation schemes. Supplementary Table S1 shows the timepoints removed in each task. We consider the left-out timepoints to be the testing set, and the remaining timepoints are used to train the model. In each case, HVGs are selected based on cells corresponding to the training timepoints in order to avoid data leakage.

#### Baselines

We compare `scNODE` with two main state-of-the-art generative methods that are capable of both interpolating and extrapolating gene expression:

- **PRESCIENT**: Yeo et al. (2021) propose a generative model, called Potential eneRgy undErlying Single Cell gradIENTs (`PRESCIENT`), to learn the differentiation landscape from single-cell time-series gene expression data. `PRESCIENT` maps single-cell gene expression to a lower-dimensional PCA space and models cell differentiation with a neural ODE.
- **MIOFlow**: Huguet et al. (2022) integrate geodesic VAE and neural ODE to model the paths of cells in a lower-dimensional latent space while cellular variations are preserved.

#### Hyperparameter tuning

In each task for every dataset, we select corresponding hyperparameters for all methods (our and baselines) that yield the minimum averaged Wasserstein distance using the 3-fold cross-validation scheme. We use Optuna (Akiba et al., 2019) to automatically determine the optimal hyperparameters and use sufficiently large hyperparameter ranges for search and evaluation. The hyperparameter ranges of `scNODE` and baselines are listed in Supplementary Table S2. We set the latent space dimension as 50, use the first-order Euler ODE solver, and set ODE step size Δ*t* = 0.1 for all methods, and run each method for sufficient iterations to ensure they converge. We let every model predict 2000 cells at each testing timepoint to ensure a fair metric comparison. Moreover, we evaluate `scNODE` performance using different hyperparameter settings, conduct ablation studies, and give heuristic guidance on how to set hyperparameters in real-world scenarios in Supplementary Sec. S6.3.

### 3.2 scNODE **can accurately predict gene expression at unobserved timepoints**

We compare `scNODE` ‘s performance with baseline methods for predicting gene expression for left-out timepoints. Fig. 2 (along with Supplementary Fig. S1∼S3) visualizes true gene expression values and model predictions for left-out testing timepoints in 2D Uniform Manifold Approximation and Projection (UMAP) (McInnes et al., 2018) space. Our results indicate `scNODE` ‘s predictions align well with ground truth. Qualitative evaluation of all scRNA-seq datasets indicates that `scNODE` can accurately predict gene expression at missing timepoints.

**Figure 2:**
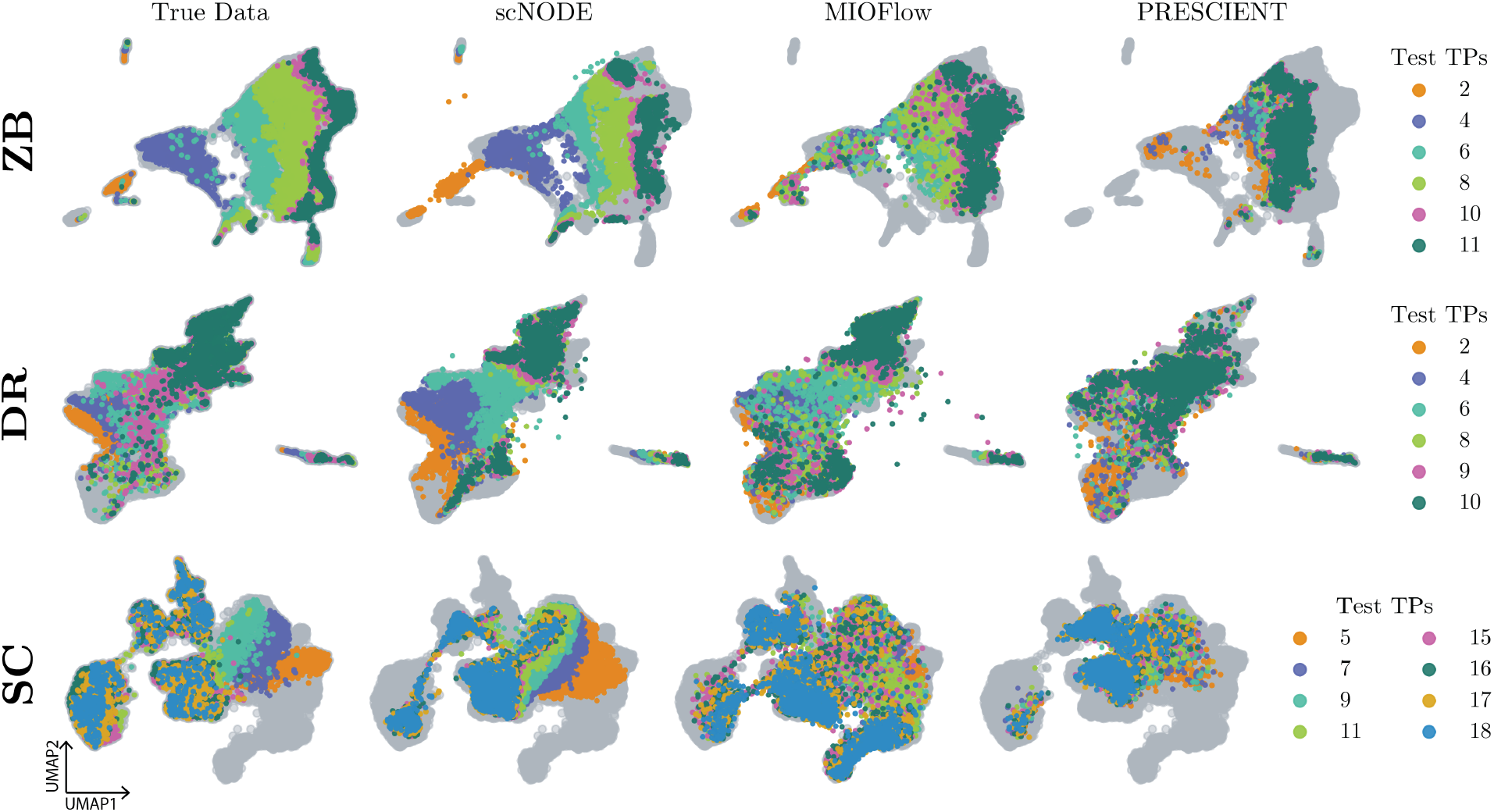
2D UMAP visualization of true and predicted gene expressions in hard tasks. Gray points represent training data.

We then quantitatively evaluate model predictions. We use the Wasserstein distance to measure the similarity between true and predicted gene expression at each testing timepoint, where a lower value corresponds to more accurate predictions. This evaluation metric has been used in previous studies (Yeo et al., 2021; Tong et al., 2020) because Wasserstein distance learns a soft mapping across the distributions, and we do not have one-to-one cell matching between true and predicted gene expression for single-cell measurements. Table 2 shows the Wasserstein distance of left-out testing timepoints averaged on five trials for defined hard tasks, and Supplementary Table S3 shows metrics for easy and medium tasks. `scNODE` clearly outperforms the baselines in most cases. In other cases where `scNODE` has the second best predictions, such as *t* = 7 of SC hard task, `scNODE` has similar performance as `MIOFlow` but significantly performs better than `PRESCIENT`. Moreover, in medium and hard tasks, where extrapolations are required, `scNODE` can have substantial improvement over baselines. For example, at *t* = 15 of SC hard task (in Table 2), Wasserstein distance of `scNODE` predictions is around 132, while that of `PRESCIENT` and `MIOFlow` are about 150 and 162 respectively. In Supplementary Fig. S4, we further evaluate all model predictions when extrapolating more timepoints. As expected, their extrapolations become less accurate the farther out predictions are made. But `scNODE` still demonstrates improvement in most cases. Therefore, `scNODE` has consistent good performance, especially in more challenging prediction tasks.

**Table 2:** Wasserstein distance of predictions in hard tasks (inter- and extrapolation). **Bold** numbers denote the best prediction, and underlined numbers represent the second best.

| Method | ZB / Hard Task |  |  |  |  |  |
| --- | --- | --- | --- | --- | --- | --- |
|  | Interpolation |  |  |  | Extrapolation |  |
| | $t = 2$ | $t = 4$ | $t = 6$ | $t = 8$ | $t = 10$ | $t = 11$ |
| scNODE | **579.10** | **508.55** | **440.92** | **517.81** | **652.36** | **707.10** |
| MIOFlow | <u>580.18</u> | <u>516.59</u> | <u>453.61</u> | <u>536.35</u> | <u>671.23</u> | <u>734.42</u> |
| PRESCIENT | 1381.96 | 1002.62 | 730.974 | 701.29 | 916.51 | 973.17 |

| Method | DR / Hard Task |  |  |  |  |  |
| --- | --- | --- | --- | --- | --- | --- |
|  | Interpolation |  |  |  | Extrapolation |  |
| | $t = 2$ | $t = 4$ | $t = 6$ | $t = 8$ | $t = 9$ | $t = 10$ |
| scNODE | <u>445.82</u> | **464.78** | <u>535.78</u> | **600.18** | <u>585.60</u> | **718.20** |
| MIOFlow | **443.56** | <u>469.51</u> | **532.93** | <u>617.48</u> | 680.41 | 852.02 |
| PRESCIENT | 524.38 | 511.61 | 539.38 | 621.31 | **575.45** | <u>718.56</u> |

| Method | SC / Hard Task |  |  |  |  |  |  |  |
| --- | --- | --- | --- | --- | --- | --- | --- | --- |
|  | Interpolation |  |  |  | Extrapolation |  |  |  |
| | $t = 5$ | $t = 7$ | $t = 9$ | $t = 11$ | $t = 15$ | $t = 16$ | $t = 17$ | $t = 18$ |
| scNODE | <u>55.22</u> | **59.89** | **103.26** | **140.81** | **132.86** | **148.89** | **137.90** | **151.13** |
| MIOFlow | **55.07** | <u>61.80</u> | <u>108.72</u> | 156.51 | 162.12 | 191.40 | 189.39 | 215.74 |
| PRESCIENT | 85.36 | 87.47 | 114.16 | <u>142.03</u> | <u>150.53</u> | <u>161.59</u> | <u>147.23</u> | <u>155.06</u> |

Next, we validate that `scNODE` is more robust to distribution shift when testing timepoints have substantially different distributions from training data. Specifically, for the hard task of all three datasets, we compute the distribution shift level for each testing timepoint as the averaged pairwise *𝓁*_2_ distance between cells from training and testing timepoints. Hence, a higher value indicates the testing point has a more significant difference from the training data. Moreover, we define `scNODE` ‘s improvement as the difference between the Wasserstein distances of the predictions made by `scNODE` and the second-best baseline, such that a higher value indicates that `scNODE` obtains more improvements. Fig. 3 shows that `scNODE` improvements positively correlate with the distribution shift level (Spearman correlation *ρ* ≥ 0.3 and is as high as 0.93.). Therefore, due to the dynamic regularization introduced in our framework, `scNODE` is more robust to the distribution shift between the observed and unobserved timepoints and obtains significant improvements over baseline models.

**Figure 3:**
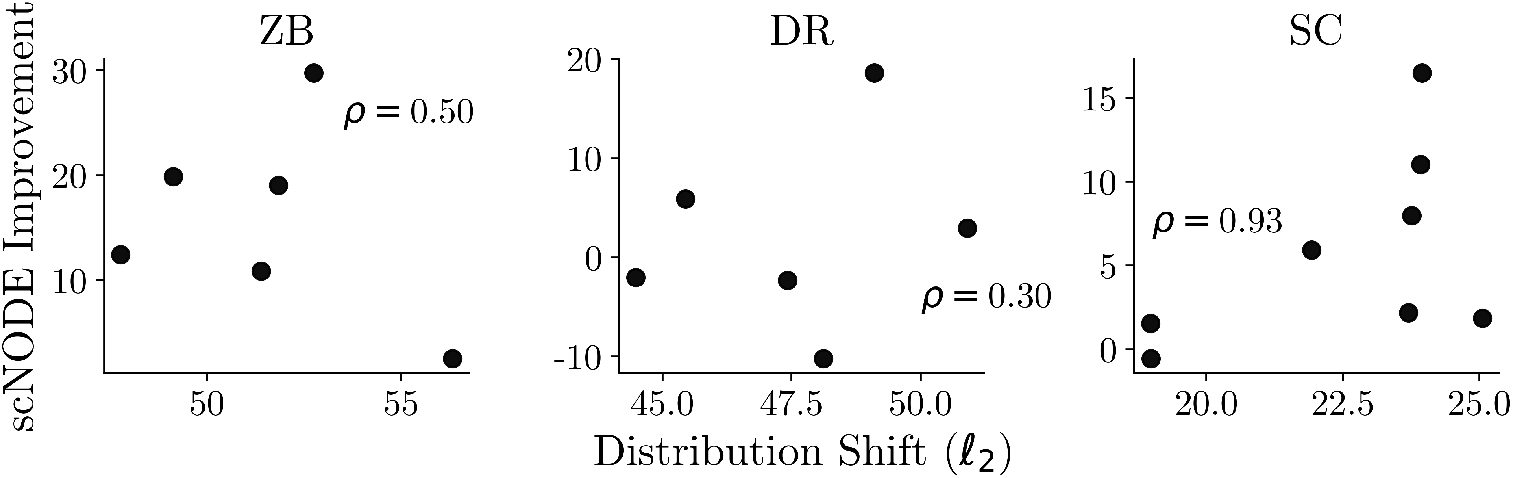
`scNODE` improvements over baseline models vs. distribution shift levels of testing points on hard tasks. Each point denotes one testing timepoint with `scNODE` improvements averaged on five trials. *ρ*: Spearman’s *ρ* correlations between improvements and distribution shift.

### 3.3 scNODE **predictions for unobserved timepoints help recover cell trajectories**

To determine if `scNODE` can aid with temporal downstream analysis, we carry out trajectory analysis on both real and predicted single-cell gene expression. Specifically, we focus on the hard task of all three datasets. Fig. 4 visualizes the ZB data with all timepoints, after the removal of testing timepoints, and with predictions from all models. We apply partition-based graph abstraction (PAGA) (Wolf et al., 2019), a method that computes the topological structures of cellular populations and has been used in previous studies (Saelens et al., 2019; Luecken and Theis, 2019; Han et al., 2020) to understand cell developmental topologies. We find that after timepoint removal, the topology breaks down due to the gaps between timepoints, which can impede the trajectory analysis, while `scNODE` predictions recover smooth and continuous trajectories. To quantitatively compare `scNODE` with other models in helping infer cell trajectories, we use the Ipsen-Mikhailov (IM) distance (Ipsen, 2004; Saelens et al., 2019) to measure the similarity between the cell trajectory graphs constructed in each case. IM(*𝒢*_1_, *𝒢*_2_) is a graph similarity measurement defined as the square-root difference between the Laplacian spectrum of graphs *𝒢*_1_ and *𝒢*_2_. It ranges from 0 to 1, where 0 indicates maximum similarity between two graph structures and 1 indicates maximum dissimilarity. We find IM(*𝒢*_true_, *𝒢*_removal_) = 0.200 and IM(*𝒢*_true_, *𝒢*_scNODE_) = 0.093, indicating *𝒢*_scNODE_ is more similar to *𝒢*_true_ than *𝒢*_removal_ such that `scNODE` predictions help recovering cell trajectories. Moreover, IM(*𝒢*_true_, *𝒢*_scNODE_) being smaller than IM(*𝒢*_true_, *𝒢*_MIOFlow_) and IM(*𝒢*_true_, *𝒢*_PRESCIENT_) (Fig. 4) implies that `scNODE` predictions for missing timepoints best help to infer cell trajectories. We observe same trend for DR and SC datasets as well, where IM distance is lowest for `scNODE` (Supplementary Fig. S8).

**Figure 4:**
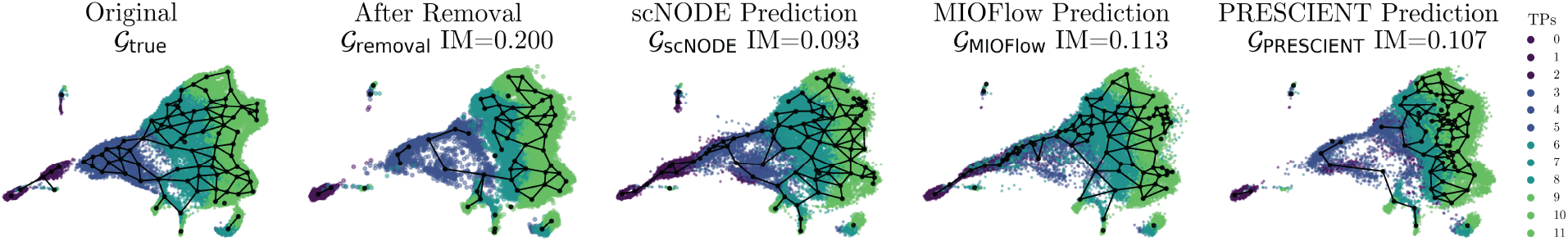
Cell trajectories of ZB data with all timepoints, after removal of timepoints in the hard task, and with model predictions. The connective structure is constructed with PAGA, where black nodes represent cell clusters and edges connect two nodes if their expressions are similar. We show the IM index between *𝒢*_true_ and the corresponding graph.

### 3.4 scNODE ‘s interpretable latent space assists with perturbation analysis

`scNODE` ‘s ability to learn an informative and robust latent space allows it to detect key driver genes for cell developmental paths and simulate cells with perturbed gene expression. This ability can be useful for performing *in silico* perturbation predictions. We validate this hypothesis by using the differential dynamics learned from the ZB dataset. We choose ZB data because the original dataset provides cell type labels at the last timepoint, which enables us to evaluate the perturbation predictions.

We first train `scNODE` with all timepoints in the ZB dataset and map all cells to the latent space with the learned `scNODE` encoder. In the latent space, we construct a most probable path between any two points through the Least Action Path (LAP) method (Qiu et al., 2022, 2012; Wang et al., 2014), which finds the optimal path between two cell states while minimizing its action and transition time (see Supplementary Sec. S6.5). Here, we construct LAP paths from cells at the starting point (i.e., *t* = 0) to presomitic mesoderm (PSM) and Hindbrain cell populations (Fig. 5**A**)

**Figure 5:**
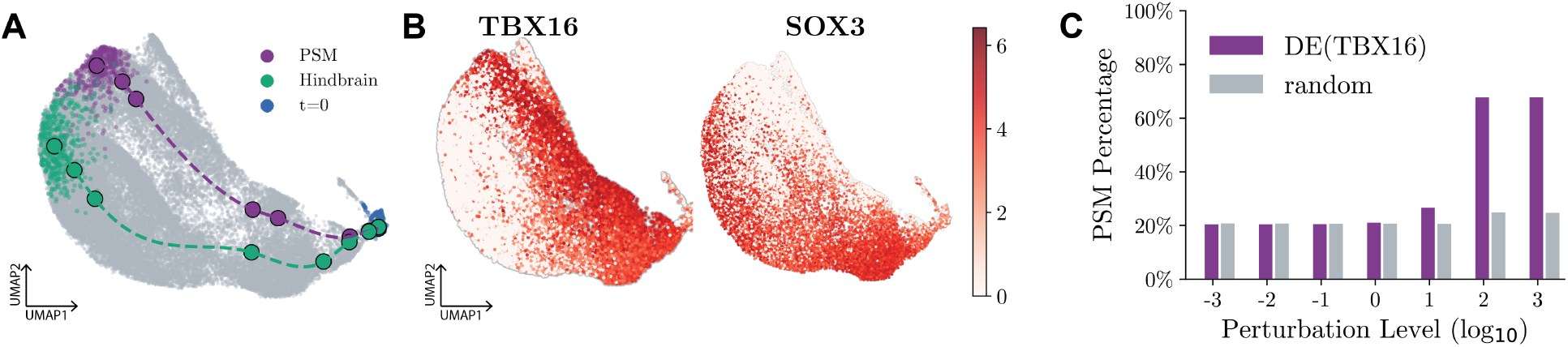
`scNODE` perturbation analysis results. (**A**) 2D UMAP visualization of the least action path between cells at the starting point (*t* = 0) and the PSM and Hindbrain cell populations, respectively. (**B**) Gene expression value of the top-rank DE gene of PSM path (TBX16) and Hindbrain path (SOX3). (**C**) The ratio of PSM cells in predictions of different perturbation levels when perturbing DE genes and random non-DE genes.

Once the optimal path is constructed, we use the Wilcoxon rank-sum test to find differentially expressed (DE) genes for each path, representing key driver genes for the developmental trajectories. Fig. 5 shows the expression of top-rank DE genes for the PSM path (TBX16) and Hindbrain path (SOX3), such that their respective expression levels vary along the path (as well as in adjacent regions). Warga et al. (2013) have found TBX16 regulates intermediate mesoderm cell fate. Other studies (Dee et al., 2008; Vriz et al., 1996) have found SOX3 regulates neural fates and is involved in central nervous system development.

Finally, we perturb the expression profiles of key genes in all cells at the starting timepoint by multiplying their expression values with different levels of coefficient *{*10^≥3^, …, 10^3^*}* to mimic overexpressing and knocking-out, and let `scNODE` predict trajectories for the perturbed gene expression. The perturbations are expected to result in changes in cell fates. We classify predicted cells with a Random Forest (RF) classifier (Breiman, 2001) trained with all unperturbed cells. We find overexpressing TBX16 expression results in the increment of PSM cell ratios (Fig. 5**C**) to over 70%, while perturbing random non-DE genes leads to no changes in cell ratios. Moreover, the ratio of Hindbrain cells decreased by around 25% when we overexpressed SOX3 expressions (Supplementary Fig. S9). These results indicate that `scNODE` learns an interpretable latent space, where we can detect development-related DE genes and conduct *in silico* perturbation analysis.

## 4 Conclusion

We propose a generative model called `scNODE` to predict gene expression during cell development. Given a time-series scRNA-seq data set, our model can both interpolate and extrapolate gene expression. Predicting gene expression for unobserved timepoints enables critical downstream analyses, such as differential gene expression analysis, perturbation analysis, and trajectory inference. To make accurate predictions, `scNODE` uses a non-linear dimensionality reduction approach, neural ODE, and dynamic regularization. The regularization improves the robustness of the learned latent representation to distribution shifts in unobserved timepoints, resulting in better predictive performance.

We find that the primary challenge for the temporal scRNA-seq prediction task is extrapolating to multiple timepoints beyond the last observed timepoint, as the accuracy deteriorates the farther timepoint predictions. Moreover, time units in predictions are defined by time intervals in the data. For example, a generative model cannot accurately produce gene expression for every hour if the model is trained on observations that are only measured every day, especially when the development is rapid or contains complex trajectories.

In future work, we will incorporate knowledge about cell proliferation - which is known to be important to cellular differentiation - into the model to improve extrapolating performance. Also, although RNA velocity methods can be used as complementary approaches to `scNODE` to learn fine-grained dynamics from coarse-grained timepoints. We will also incorporate other modalities, such as single-cell chromatin accessibility (scATAC) datasets, which exhibit a regulatory mechanism that scRNA-seq data might not capture.

## S1 Related Works

One category of methods learns gene expression dynamics by inferring cell trajectories (Saelens et al., 2019; Herman et al., 2018; Wolf et al., 2019; Trapnell et al., 2014; Qiu et al., 2017). However, most of them are based on pseudo-time inference, which estimates differential time for cells based on cell pair distances in low-dimensional spaces. Therefore, they cannot model developmental dynamics with respect to real physical time.

In addition, a category of approaches uses RNA velocity (La Manno et al., 2018; Bergen et al., 2021, 2020) to uncover developmental trends and underlying kinetics of gene expression. But they have inevitable limitations on gene expression prediction because they use linear ODEs (Matsumoto et al., 2017) that fail to capture complex gene expression dynamics, deeply sequenced datasets to obtain spliced and unspliced counts as prior information (Farrell et al., 2023), or have to use RNA velocity pre-computed from gene expression (Chen et al., 2022b) that causes double dipping (Ball et al., 2020) problems. Furthermore, most RNA velocity models are non-generative and infeasible for predicting gene expression.

Single-cell gene expression prediction has been investigated in previous studies (Cao et al., 2021). For example, Marouf et al. (2020) proposes the use of generative adversarial networks (GAN) to generate realistic augmentations for specific cell types. Moreover, Zappia et al. (2017); Risso et al. (2018); Li and Li (2019); Sun et al. (2021) use statistical models to fit distributions of observed single-cell RNA sequencing (scRNA-seq) data and use the learned information to generate simulations. These models can make accurate predictions that reflect the properties of the experimental data. However, they are designed for gene expression measured at a single time snapshot and do not consider developmental dynamics. Therefore, they are infeasible for predicting temporal scRNA-seq data.

Recently, a few generative methods have introduced procedures to understand cell differentiation based on physical time and predict unobserved timepoints. Sagittarius method (Woicik et al., 2023) predicts gene expressions at future timepoints but mainly focuses on bulk RNA-seq data that requires cell matching between timepoints. This matching is hard to obtain for real-world scRNA-seq as the cells are lysed during the experiment. Although Live-seq (Chen et al., 2022a) enables temporal transcriptomic recording on the same cells, most existing single-cell datasets still have the cell lysing problem. In another line of works, Waddington-OT (WOT) (Schiebinger et al., 2019) interpolates between two timepoints by using an optimal transport (OT) framework to predict cell-cell couplings. However, it cannot extrapolate to future timepoints as the cell-cell coupling is unavailable when the subsequent timepoint is missing. TrajectoryNet (Tong et al., 2020) utilizes continuous normalizing flow to model the paths between cell populations of two time snapshots and predict interpolations. Therefore, WOT and TrajectoryNet are unable to extrapolate beyond the observed timepoints because they require gene expression measurements at two endpoints to learn intermediate cellular developments. In addition, Sha et al. (2024) jointly models developmental dynamics with cell growth. However, it focuses on reconstructing cellular trajectories and inferring underlying gene regulatory networks. It does not show abilities to predict extrapolation to timepoints beyond the measured time range.

PRESCIENT (Yeo et al., 2021) applies principal component analysis (PCA) to reduce high-dimensional gene space to a low-dimensional representation. It then applies neural ordinary differential equations (ODEs) (Chen et al., 2018) in this low-dimensional space to model the cell developmental trajectory. Multiple studies (Ding et al., 2018; Xiang et al., 2021; Tran et al., 2021) have found that PCA-based low-dimensional representations of scRNA-seq expression cannot capture complex relationships in the highly heterogeneous single-cell data. Thus, PCA may have the issue of overcrowding representation (Tran et al., 2021; Kobak and Berens, 2019), where cells of different types are poorly separated in PCA-based low-dimensional space, and cellular variations are lost. MIOFlow (Huguet et al., 2022) replaces PCA with a geodesic variational autoencoder (VAE) to learn non-linear low-dimensional representations that better retain cellular variations. However, this method fixes its low-dimensional space at the beginning, followed by neural ODE modeling. We hypothesize that a fixed low-dimensional representation obtained using observed timepoints may not generalize to an unobserved timepoint, if its underlying distribution differs substantially. Therefore, these methods either use less effective dimensionality reduction approaches or freeze low-dimensional representations so that they have inaccurate predictions at unmeasured timepoints.

On the contrary, our `scNODE` uses variational autoencoder (VAE), a non-linear dimensionality reduction method that captures complex cell relationships. `scNODE` also updates the low-dimensional latent space when learning developmental dynamics, such that the latent space considers overall developments and is more robust against distribution shifts.

## S2 Single-Cell Dataset and Pre-Processing

We use three publicly available single-cell RNA sequencing (scRNA-seq) datasets. The ZB dataset is downloaded from Single Cell Portal with identifier SCP126. The DR data is downloaded from https://shendure-web.gs.washington.edu/content/members/DEAP_website/public/ and SC data are available at https://broadinstitute.github.io/wot/tutorial/. For all datasets, we use data after removing batch effects among different timepoints.

For each task, we first select highly variable genes (HVGs). HVGs are a subset of genes that contribute strongly to cell-to-cell variation within a cell population. Using only HVGs for analysis helps remove data noise and reduce computational costs. In each case, we detect the top 2000 HVGs only from training timepoints to avoid data leakage. We use *Scanpy* (Wolf et al., 2018) and *Serurat* (Hao et al., 2021) to detect HVGs.

We then normalize expression to remove cell-specific bias before conducting experiments. Specifically, given the unique molecular identifier (UMI) count expression of cell *i* as 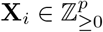 where ℤ_≥0_ = {0, 1, 2, … }, we normalize it by total counts over all genes

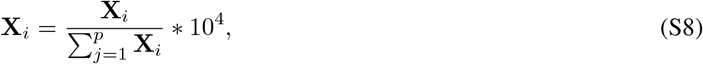

followed by the log-transformation

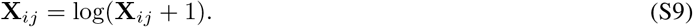

The normalization is cell-specific so that there is no data leakage between training and testing data. Because SC datasets only provide normalized expression data, we normalize the other two datasets.

**Table S1:** The leave-out timepoints (testing timepoints) for three defined tasks in each dataset. The timepoint index starts with 0. Since the SC dataset has more timepoints, we leave out more timepoints than other datasets in its hard task.

| Dataset | # Timepoints | Leave-Out Timepoints<br>(Testing Timepoints) |  |  |
| --- | --- | --- | --- | --- |
|  |  | Easy Task | Medium Task | Hard Task |
| **ZB** | 12 | 4, 6, 8 | 10, 11 | 2,4,6,8,10,11 |
| **DR** | 11 | 4, 6, 8 | 8, 9, 10 | 2,4,6,8,9,10 |
| **SC** | 19 | 5, 10, 15 | 16, 17, 18 | 5,7,9,11,15,16,17,18 |

## S3 scNODE Training and Inference

Our `scNODE` is implemented with *Pytorch 1*.*13* (Paszke et al., 2019) and is trained end-to-end. `scNODE` training consists of two main steps. `scNODE` first pre-trains the VAE component (encoder Enc_*ϕ*_ and decoder Dec_*θ*_) with all cells of training timepoints. We use Adam optimizer to pre-train `scNODE` with a learning rate of 0.001 and 200 iterations. Then, `scNODE` optimizes both VAE and neural ODE components by minimizing the *ℒ*_dyn_. We adopt batch training and use Adam optimizer to train `scNODE` with a learning rate of 0.001 and 1000 iterations. At each training iteration, we randomly select 32 cells at *t* = 0 as a batch and predict for every training timepoint *t* ∈ *𝒯*. Because the Wasserstein distance computation is expensive, batch training improves training efficiency and enables `scNODE` usage on large-scale datasets. We use *geomloss* (Feydy et al., 2019) to compute Wasserstein distance with blur = 0.05 and scaling = 0.5. Pseudo-codes of `scNODE` are provided in Algorithm S1.

Specifically, `scNODE` training loss function (Eq. 7 in main manuscript) contains a dynamic regularization term to pose cellular dynamics to the latent space and combines structure relation and temporal development of cells. This leads to an informative, interpretable, and robust latent space. At each training timepoint *t* ∈ *𝒯*, we use the encoder to compute latent variables 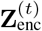 for **X**^(*t*)^ and then compute the Wasserstein distance between it and latent variables **Z**^(*t*)^ generated from neural ODE. Notice at the first timepoint *t* = 0, its latent variable **Z**^(0)^ is computed from VAE encoding, such that the regularization Wass 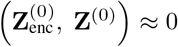. But for the latter training timepoints, this regularization enforces similarities between structure latent representations from VAE and the dynamic manifold from neural ODE.

At inference, `scNODE` first uses the trained encoder Enc_*ϕ*_ to map gene expression at the first timepoint to the latent space and then samples latent variables for the required number of cells from the latent Gaussian distribution. Then, `scNODE` uses the ODE solver to predict latent variables of these cells at required timepoints and map them back to the gene space through the decoder Dec_*θ*_. Our `scNODE` can predict gene expression at any timepoints, including those observed and unobserved ones.

## S4 Baseline Models

We compare `scNODE` with the following baseline models.

### Algorithm S1

scNODE

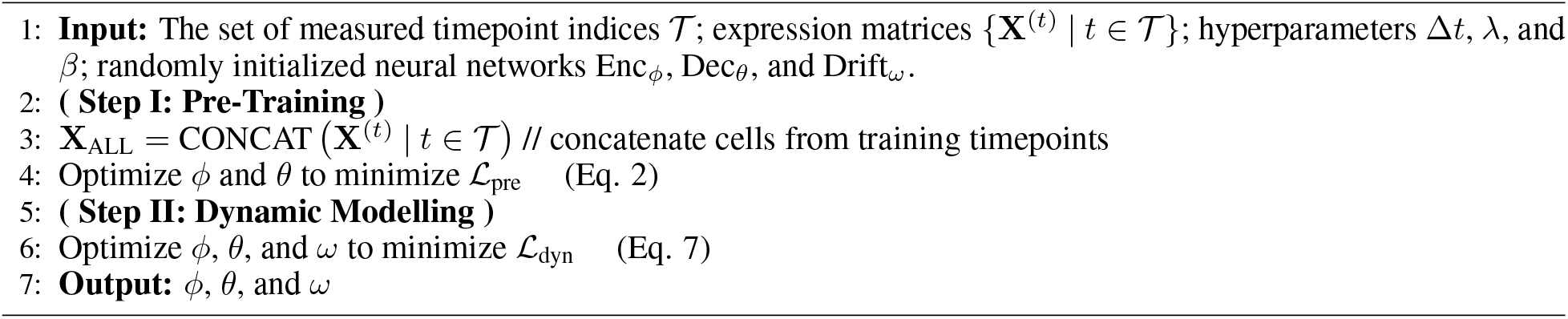

- **PRESCIENT**: Yeo et al. (2021) proposes a generative model, called Potential eneRgy undErlying Single Cell gradIENTs (`PSCIENT`), to learn the differentiation landscape from single-cell time-series gene expression data. `PRESCIENT` maps gene expression to the PCA space and models cellular differentiation with neural ordinary differential equation (ODE). `PRESCIENT` suggests using cell proliferation information to improve prediction. But cell proliferation is not available in general cases. Therefore, we do not use such information in our experiments for all methods. We use Python codes on GitHub (https://github.com/gifford-lab/ prescient) to run `PRESCIENT`.
- **MIOFlow**: Huguet et al. (2022) integrate geodesic VAE and neural ODE to model the paths of cells in a latent space while cellular structures are preserved. Specifically, `MIOFlow` first learns low-dimensional cell representation with a geodesic VAE such that latent space keeps the geometry of data manifold. `MIOFlow` then models cellular differentiation with neural ODE in this latent space. We use Python codes on GitHub (https://github.com/KrishnaswamyLab/MIOFlow) to run `MIOFlow`.

Waddintong-OT (Schiebinger et al., 2019) and TrajectoryNet (Tong et al., 2020) are the other two generative methods. However, they can only predict interpolations between timepoints. Moreover, Yeo et al. (2021) and Huguet et al. (2022) have shown that PRESCIENT and MIOFlow perform better than these two methods. In this work, we focus on generative models that can both interpolate and extrapolate gene expressions, thus excluding Waddington-OT and TrajectoryNet in the comparison.

## S5 Hyper-Parameter Tuning

For each method, we select hyperparameters that yield the minimum averaged Wasserstein distance using 3-fold cross-validation. For example, if the dataset has training timepoints *t* = 0, 1, 2 and we want to predict gene expression at testing timepoint *t* = 3. In cross-validation, we equally split the cells at every timepoint of *t* = 0, 1, 2 into three subsets, train the model on two of them, and validate it on the rest one. We repeat this process three times to ensure each subset is used as the validation set once and record the averaged Wasserstein distance on three validation sets. We use Optuna Akiba et al. (2019) to automatically determine the optimal hyperparameters and use sufficiently large hyperparameter ranges for search and evaluation. The search spaces of hyperparameters are shown in Table S2.

Specifically, we set the latent space dimension as 50 and step size Δ*t* = 0.1 for our `scNODE` and all baseline methods. State-of-the-art methods (Townes et al., 2019; Tran et al., 2021; Heumos et al., 2023) also generally choose a latent space of tens of dimensions. For a fair comparison, we set the same latent space size for all methods. Moreover, step size Δ*t* = 0.1 implies the neural ODE interpolates 10 unformaly spaced timepoints between two consecutive training timepoints. A small step size increases dynamic resolution and a larger step size reduces computational costs. Previous works (Yeo et al., 2021; Huguet et al., 2022) also use Δ*t* = 0.1 as a proper tradeoff between efficiency and accuracy. We train each method for sufficient iterations to ensure they converge. Specifically, we train `scNODE` 200 pre-training iterations and 1000 training iterations, `PRESCIENT` 500 epochs, `MIOFlow` 1000 pre-training iterations and 100 training epochs. We let every model predict 2000 cells at each predicted timepoints to ensure a fair metric comparison. The datasets used in our experiments have thousands of cells at each timepoint, so we believe 2000 is a reasonable number of predicted cells.

**Table S2:**
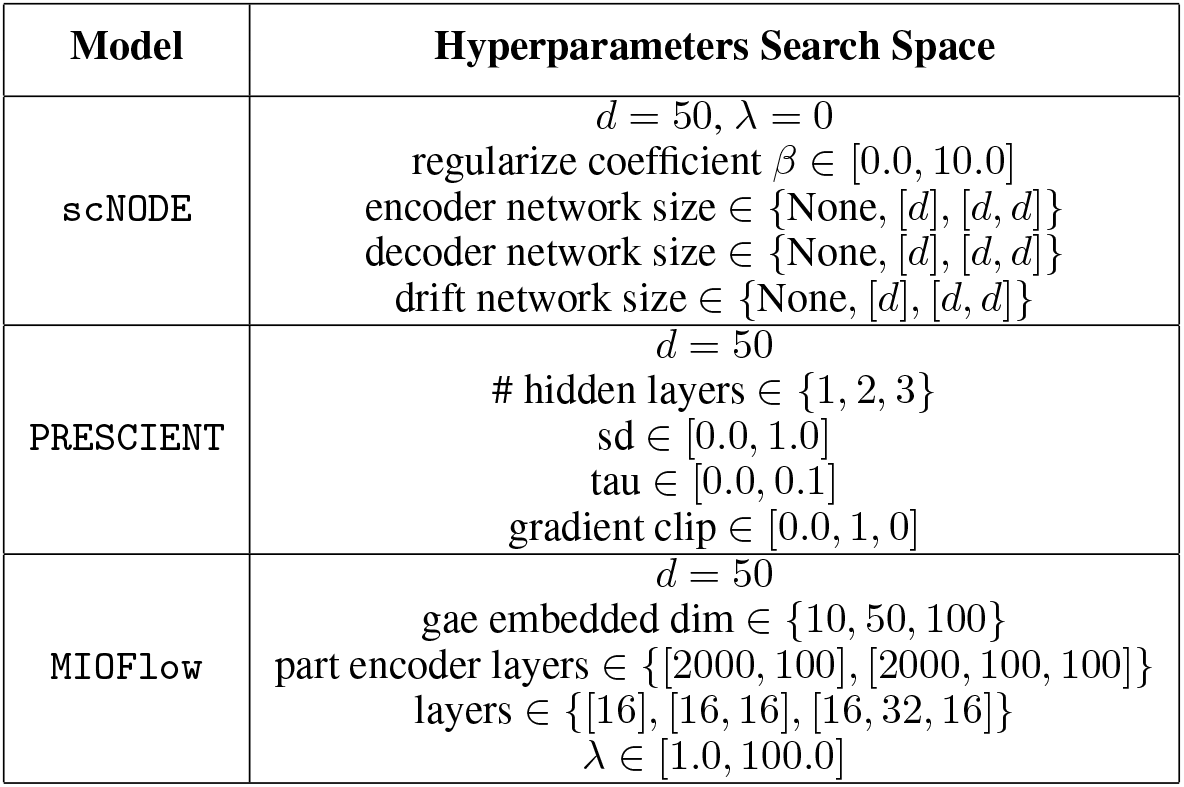
Hyperparameter search space of `scNODE` and baseline methods. “None” means no hidden layers.

| Model | Hyperparameters Search Space |
| --- | --- |
| scNODE | $d = 50, \lambda = 0$<br>regularize coefficient $\beta \in [0.0, 10.0]$<br>encoder network size $\in \{\text{None}, [d], [d, d]\}$<br>decoder network size $\in \{\text{None}, [d], [d, d]\}$<br>drift network size $\in \{\text{None}, [d], [d, d]\}$ |
| PRESCIENT | $d = 50$<br># hidden layers $\in \{1, 2, 3\}$<br>sd $\in [0.0, 1.0]$<br>tau $\in [0.0, 0.1]$<br>gradient clip $\in [0.0, 1, 0]$ |
| MIOFlow | $d = 50$<br>gae embedded dim $\in \{10, 50, 100\}$<br>part encoder layers $\in \{[2000, 100], [2000, 100, 100]\}$<br>layers $\in \{[16], [16, 16], [16, 32, 16]\}$<br>$\lambda \in [1.0, 100.0]$ |

**Table S3:**
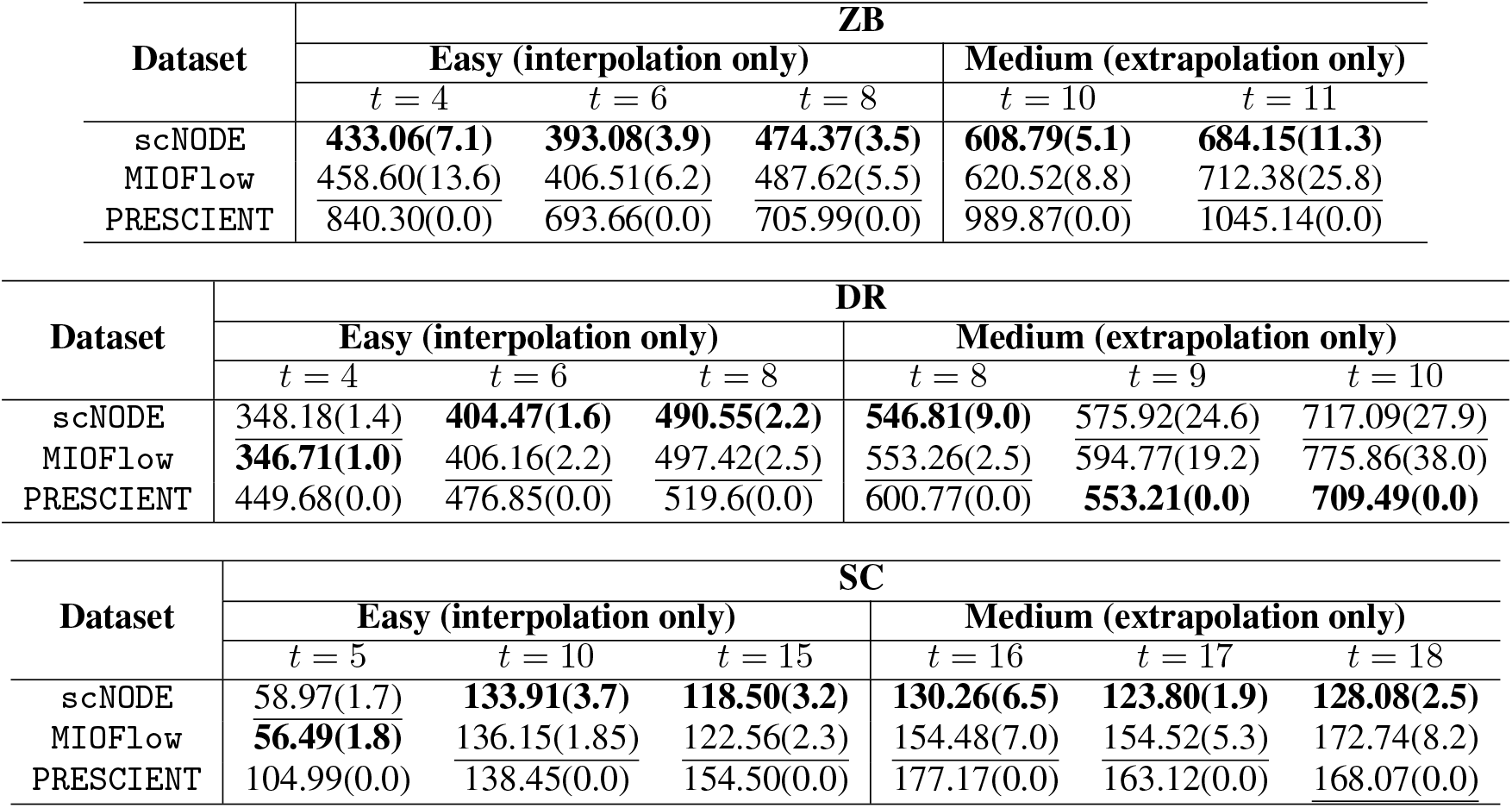
Wasserstein distance of predictions in the easy (interpolation only) and medium (extrapolation only) tasks for all three real-world datasets. We report average(std.) Wasserstein distance over five trials. Bold numbers denote the best prediction, and underlined numbers represent the second best.

| Dataset | ZB |  |  |  |  |  |
| --- | --- | --- | --- | --- | --- | --- |
|  | Easy (interpolation only) |  |  | Medium (extrapolation only) |  |  |
| | $t = 4$ | $t = 6$ | $t = 8$ | $t = 10$ | $t = 11$ | |
| scNODE | **433.06(7.1)** | **393.08(3.9)** | **474.37(3.5)** | **608.79(5.1)** | **684.15(11.3)** |  |
| MIOFlow | 458.60(13.6) | 406.51(6.2) | 487.62(5.5) | 620.52(8.8) | 712.38(25.8) |  |
| PRESCIENT | 840.30(0.0) | 693.66(0.0) | 705.99(0.0) | 989.87(0.0) | 1045.14(0.0) |  |

| Dataset | DR |  |  |  |  |  |
| --- | --- | --- | --- | --- | --- | --- |
|  | Easy (interpolation only) |  |  | Medium (extrapolation only) |  |  |
| | $t = 4$ | $t = 6$ | $t = 8$ | $t = 8$ | $t = 9$ | $t = 10$ |
| scNODE | 348.18(1.4) | **404.47(1.6)** | **490.55(2.2)** | **546.81(9.0)** | 575.92(24.6) | 717.09(27.9) |
| MIOFlow | **346.71(1.0)** | 406.16(2.2) | 497.42(2.5) | 553.26(2.5) | 594.77(19.2) | 775.86(38.0) |
| PRESCIENT | 449.68(0.0) | 476.85(0.0) | 519.6(0.0) | 600.77(0.0) | **553.21(0.0)** | **709.49(0.0)** |

| Dataset | SC |  |  |  |  |  |
| --- | --- | --- | --- | --- | --- | --- |
|  | Easy (interpolation only) |  |  | Medium (extrapolation only) |  |  |
| | $t = 5$ | $t = 10$ | $t = 15$ | $t = 16$ | $t = 17$ | $t = 18$ |
| scNODE | 58.97(1.7) | **133.91(3.7)** | **118.50(3.2)** | **130.26(6.5)** | **123.80(1.9)** | **128.08(2.5)** |
| MIOFlow | **56.49(1.8)** | 136.15(1.85) | 122.56(2.3) | 154.48(7.0) | 154.52(5.3) | 172.74(8.2) |
| PRESCIENT | 104.99(0.0) | 138.45(0.0) | 154.50(0.0) | 177.17(0.0) | 163.12(0.0) | 168.07(0.0) |

## S6 More on experiments

### S6.1 scNODE can accurately predict expression at unmeasured timepoints

We compare `scNODE` ‘s performance with baseline methods for predicting gene expression at left-out timepoints. Tables S3 and S4 show the Wasserstein distance of left-out testing timepoints for defined easy, medium, and hard tasks. We report Wasserstein distances averaged on five repeating trials. `scNODE` clearly outperforms the baselines in most cases. In other cases where `scNODE` has the second best predictions, `scNODE` has a similar performance as `MIOFlow` and significantly performs better than `PRESCIENT`. Moreover, in medium and hard tasks, where extrapolations are required, `scNODE` can have significant superiority over baselines. For example, at *t* = 15 of SC hard task (in Table S4), Wasserstein distance of `scNODE` predictions is around 132, while that of `PRESCIENT` and `MIOFlow` are about 162 and 150 respectively. Therefore, `scNODE` has better overall performance, especially in more challenging tasks.

**Table S4:** Wasserstein distance of predictions in hard tasks (inter- and extrapolation). We report average(std.) Wasserstein distance over five trials. **Bold** numbers denote the best prediction, and underlined numbers represent the second best.

| Method | ZB / Hard Task |  |  |  |  |  |
| --- | --- | --- | --- | --- | --- | --- |
|  | Interpolation |  |  |  | Extrapolation |  |
| | $t = 2$ | $t = 4$ | $t = 6$ | $t = 8$ | $t = 10$ | $t = 11$ |
| scNODE | **579.10(2.2)** | **508.55(3.6)** | **440.92(4.7)** | **517.81(2.2)** | **652.36(3.1)** | **707.10(4.9)** |
| MIOFlow | 580.18(7.5) | 516.59(7.9) | 453.61(3.4) | 536.35(2.3) | 671.23(15.9) | 734.42(21.7) |
| PRESCIENT | 1381.96(0.0) | 1002.62(0.0) | 730.974(0.0) | 701.29(0.0) | 916.51(0.0) | 973.17(0.0) |

| Method | DR / Hard Task |  |  |  |  |  |
| --- | --- | --- | --- | --- | --- | --- |
|  | Interpolation |  |  |  | Extrapolation |  |
| | $t = 2$ | $t = 4$ | $t = 6$ | $t = 8$ | $t = 9$ | $t = 10$ |
| scNODE | 445.82(4.7) | **464.78(2.6)** | 535.78(3.8) | **600.18(7.0)** | 585.60(6.8) | **718.20(3.2)** |
| MIOFlow | **443.56(6.5)** | 469.51(1.7) | **532.93(2.5)** | 617.48(9.8) | 680.41(18.6) | 852.02(37.2) |
| PRESCIENT | 524.38(0.0) | 511.61(0.0) | 539.38(0.0) | 621.31(0.0) | **575.45(0.0)** | 718.56(0.0) |

| Method | SC / Hard Task |  |  |  |  |  |  |  |
| --- | --- | --- | --- | --- | --- | --- | --- | --- |
|  | Interpolation |  |  |  | Extrapolation |  |  |  |
| | $t = 5$ | $t = 7$ | $t = 9$ | $t = 11$ | $t = 15$ | $t = 16$ | $t = 17$ | $t = 18$ |
| scNODE | 55.22(0.9) | **59.89(1.0)** | **103.26(2.0)** | **140.81(2.4)** | **132.86(7.3)** | **148.89(10.8)** | **137.90(10.6)** | **151.13(12.7)** |
| MIOFlow | **55.07(2.0)** | 61.80(1.6) | 108.72(2.1) | 156.51(6.9) | 162.12(19.1) | 191.40(30.2) | 189.39(35.3) | 215.74(49.9) |
| PRESCIENT | 85.36(0.0) | 87.47(0.0) | 114.16(0.0) | 142.03(0.0) | 150.53(0.0) | 161.59(0.0) | 147.23(0.0) | 155.06(0.0) |

We also visualize ground truth and model predictions in the 2D UMAP space to qualitatively compare `scNODE` with baseline models. Specifically, in each case, we first map all timepoints of ground truth to the 50-dimensional PCA space and then compute the 2D UMAP embedding of them. Then we project each model’s predictions to this UMAP space with the PCA and UMAP transformation initialized from ground truth. We use the *umap* package (https://github.com/lmcinnes/umap) and set n_neighbors=50 and min_dist=0.1 when computing UMAP embeddings. Fig. S1, S2, and S3 show that our `scNODE` predictions better align with ground truth.

**Figure S1:**
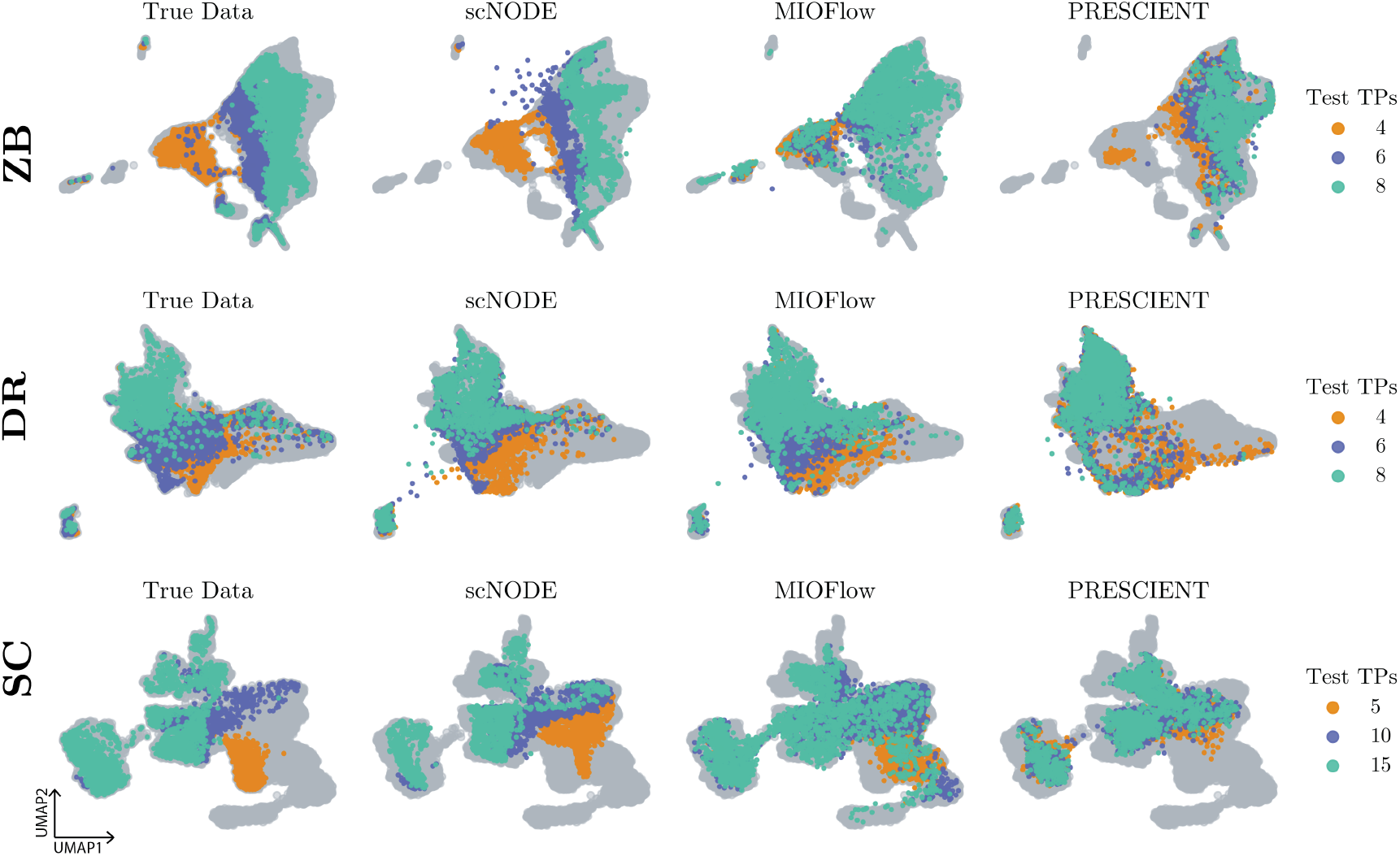
UMAP visualizations of true and model predictions in easy tasks. Gray points represent training data.

Unsurprisingly, extrapolating into the future is more difficult than interpolating because the training data may not contain sufficient information about future developments. Moreover, the distribution of gene expression at future timepoints may shift drastically from training data, so extrapolation is challenging for model generalization. Therefore, to evaluate model predictions in extrapolating multiple timepoints and test the generalization and stability of generative models against distribution shift, we let each model predict different future timepoints and track its performance. Specifically, in the ZB, DR, and SC datasets, we remove the last *{*1, 2 …, 5*}* timepoints, train each model on the rest of the timepoints, and let them predict extrapolations. In each case, we re-do the preprocessing and re-tune all hyperparameters. Fig. S4 shows the averaged Wasserstein distance in each setting. We notice for all methods, their extrapolations become less accurate the farther out predictions are made. But `scNODE` consistently performs better in most cases. Especially when extrapolating five timepoints, `scNODE` significantly outperforms the other two methods in DR and SC datasets, which implies that `scNODE` better generalizes to unobserved future timepoints that have shifted distributions. Therefore, although accurate extrapolation is a challenge (Woicik et al., 2023) in single-cell gene expression prediction, `scNODE` still demonstrates significant improvement over state-of-the-art methods in accurately extrapolating future timepoints.

**Figure S2:**
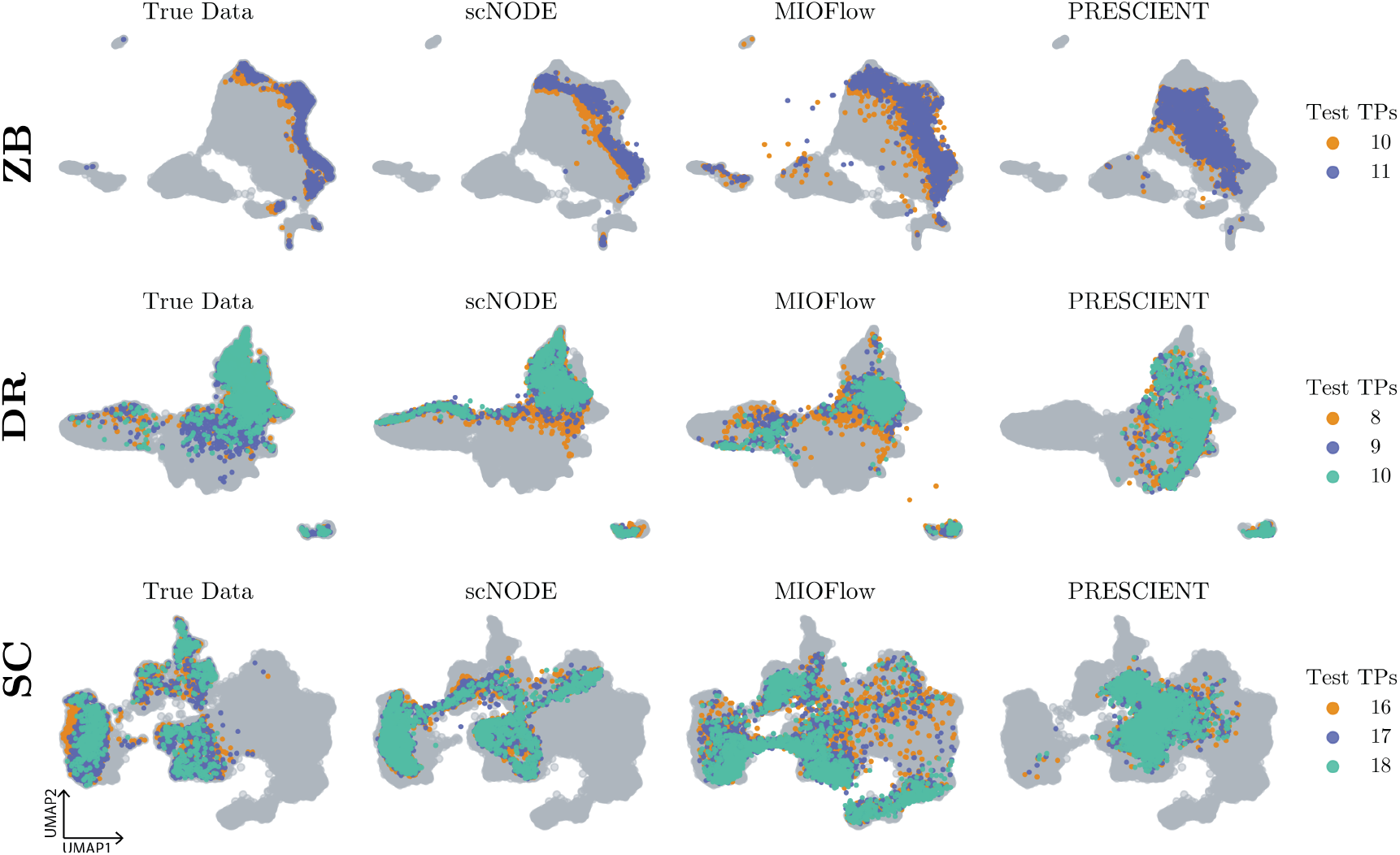
UMAP visualizations of true and model predictions in medium tasks. Gray points represent training data.

**Figure S3:**
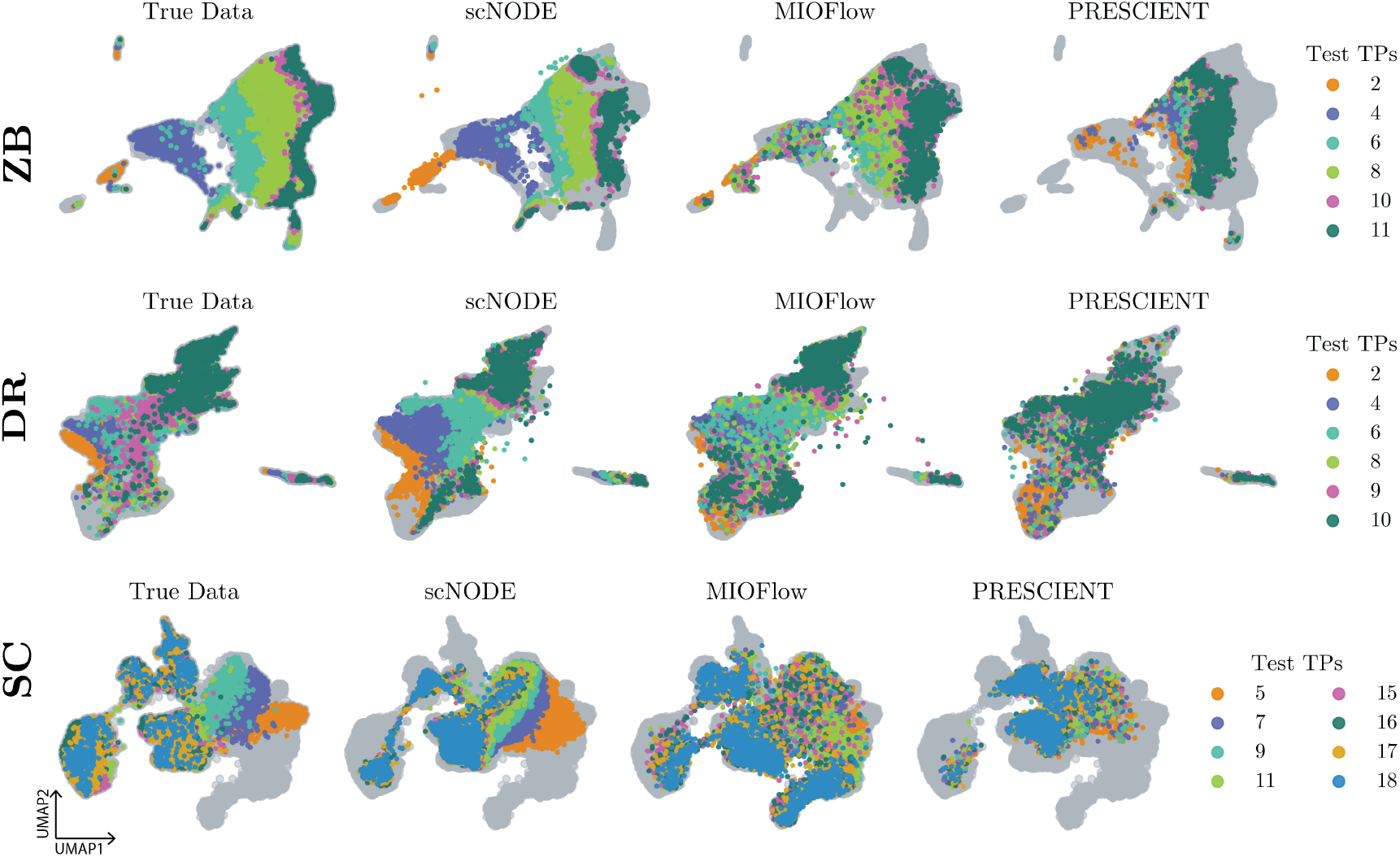
UMAP visualizations of true and model predictions in hard tasks. Gray points represent training data.

**Figure S4:**
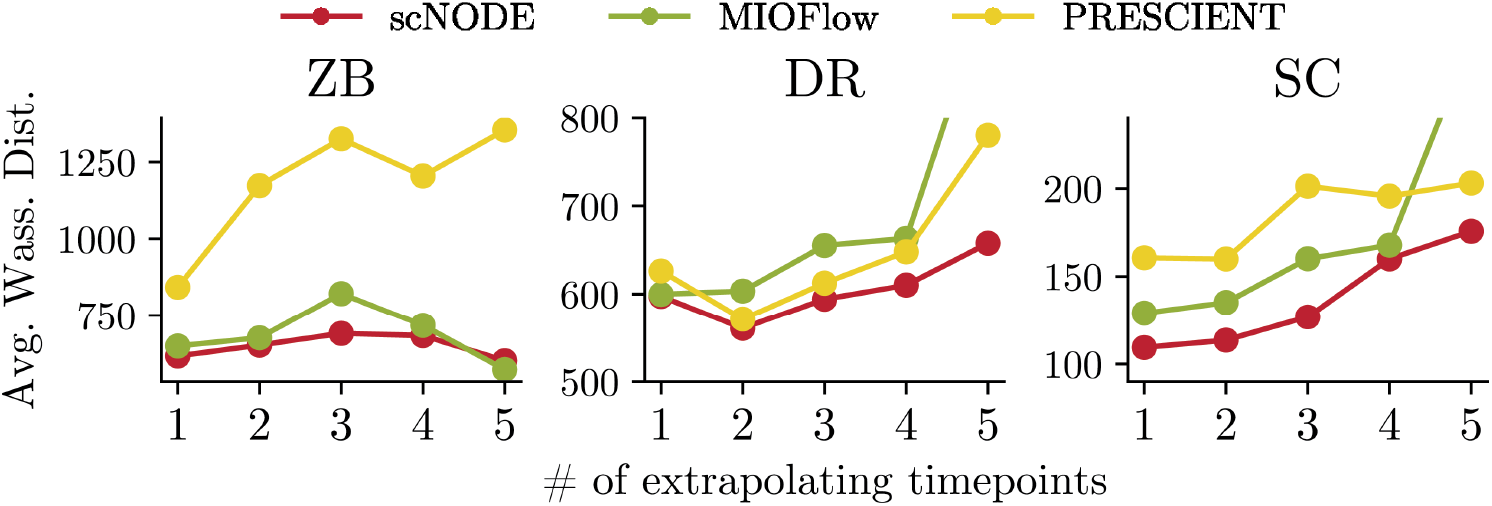
Compare `scNODE` and baseline model predictions on extrapolating multiple timepoints in ZB, DR, and SC datasets. We show the Wasserstein distance between true and predicted expression averaged across all testing timepoints.

### S6.2 scNODE is more robust to distribution shift between observed and unobserved timepoints

We show that `scNODE` is more robust to distribution shift and obtains significant improvements over baseline models when testing timepoints have substantially different distributions as training data. In three hard tasks, we compute the distribution shift level for each testing timepoint as the averaged pairwise *𝓁*_2_ distance between cells from training and testing timepoints. Specifically, given training data matrix **X** ∈ ℝ^*n×p*^ and testing data matrix **Y** ∈ ℝ^*m×p*^. The distribution shift level is computed through

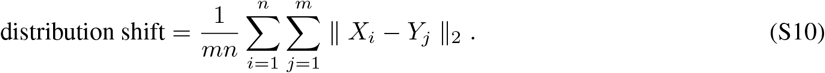

Hence, a higher value indicates the testing point has more significant differences from the training data. Moreover, we define the `scNODE` improvement as the difference between the prediction Wasserstein distance of `scNODE` and the best baseline, such that a higher value indicates that `scNODE` obtains more improvements. We compute the Spearman’s *ρ* correlation between distribution level and `scNODE` improvements for each testing timepoint. Before computing correlation, we remove outliers by fitting the data points with RANdom SAmple Consensus (RANSAC) regression, an outlier-robust regression algorithm, and remove the sample with the largest residual. Fig. 3 in the manuscript implies that `scNODE` improvements positively correlate with the distribution shift level. The correlations are *ρ* = 0.5, *ρ* = 0.3, and *ρ* = 0.93 for ZB, DR, and SC datasets, respectively. Therefore, `scNODE` is robust to distribution shift and shows more superiority when testing data are more different.

Last, we validate that `scNODE` learns a more robust latent space. We use the setting of leaving out the last three timepoints in all three datasets, where testing timepoints are located outside the training time range and have different distributions. We train all methods with training timepoints and compare the latent variables of testing timepoints generated by each model. Fig. S5 shows that `PRESCIENT` generates an over-mixing distribution of testing timepoints in the latent space, implying it fails at generating samples out of training distribution and has the worst overall performance. `scNODE` and `MIOFlow` can model temporal dynamics in the latent space and distinguish between different timepoints. However, `MIOFlow` latent variables are scattered and random, while `scNODE` has a more structured latent space, leading to accurate predictions at testing timepoints. This should be one reason for `scNODE` ‘s robustness against distribution shift.

**Figure S5:**
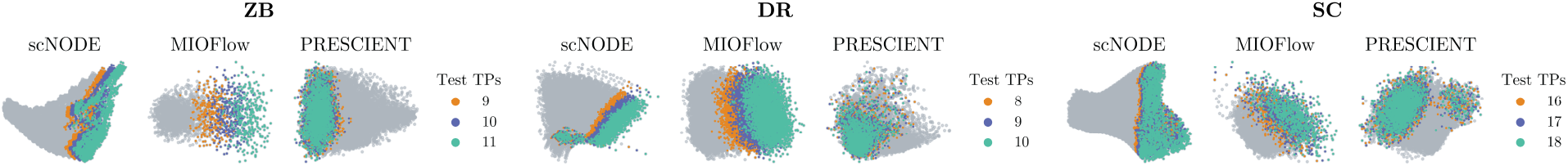
Compare two-dimensional PCA visualization of latent variables when extrapolating the last three timepoints in ZB, DR, and SC datasets.

### S6.3 Investigation of scNODE hyperparameters

We evaluate `scNODE` performance using different hyperparameter settings and give heuristic guidance on how to set hyperparameters in real-world scenarios. All experiments are conducted on the hard tasks. First, we vary the latent dimension from *{*25, 50, 75, …, 200*}*. Table S5 indicates that `scNODE` predictions are robust to the size of the latent dimensionality. Users can choose to set a reasonable latent dimension based on a tradeoff between accuracy and computational costs. State-of-the-art methods (Townes et al., 2019; Tran et al., 2021; Heumos et al., 2023) generally choose a latent space of 10 to 50 dimensions. We set the latent space size to 50 for all methods in our main comparison.

**Table S5:**
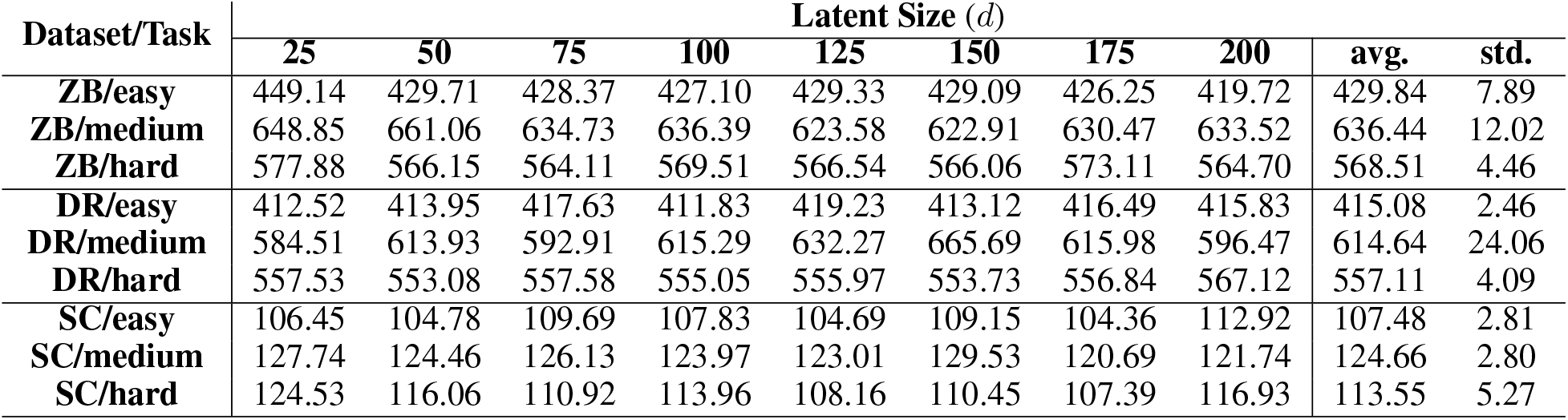
Performance of `scNODE` predictions on all datasets when using different latent space size (*d*).

Then, we test the effects of the pre-training phase. Specifically, we compare the accuracy of `scNODE` predictions when excluding the pre-training step and using different numbers of training timepoints in pre-training. We compare two strategies of choosing pretraining timepoints: randomly select from training timepoints (denoted as “random”) or pick the first several training timepoints (denoted as “first”). In each case, we run `scNODE` 5 trials. Fig. S6 shows that pre-training clearly improves predictions in all cases. Moreover, `scNODE` can outperform baselines even when not using all training timepoints in the pre-training step. This is because `scNODE` uses dynamic regularization to update the latent space. To validate the effects of latent space adjustment in `scNODE`, we also compare the predictions of `scNODE` when including and excluding the update of latent space. Excluding the adjustment of latent space means fixing the pre-trained VAE and only optimizing neural ODE parameters *ω* during model learning. Fig. S7 shows that latent space adjustment improves prediction accuracy. Overall, the ablation study shows that pre-training and latent space updates are necessary for accurate predictions.

**Figure S6:**
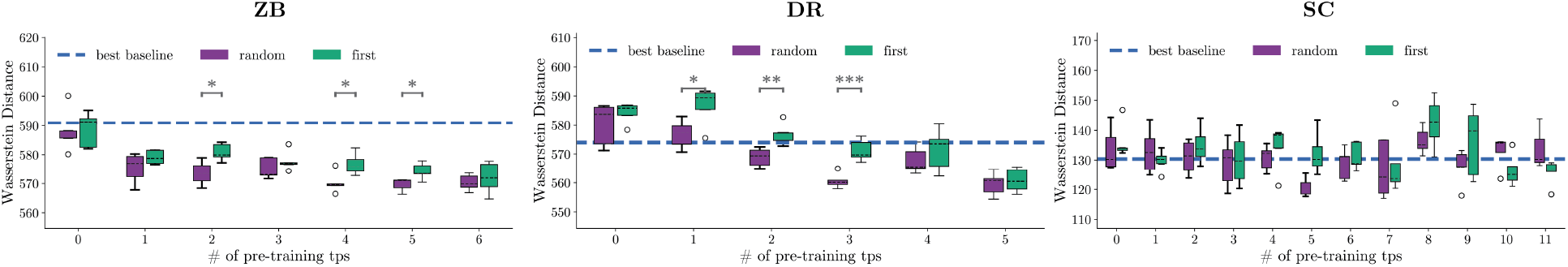
Compare Wasserstein distances of `scNODE` predictions in hard tasks when using different numbers of pre-training timepoints. Asterisks indicate p-values of the t-test between two strategies. *: p-value≥ 0.05, **: p-value≥ 0.01, and ***: p-value≥ 0.001. Dotted lines indicate the best baseline performance.

`scNODE` uses a dynamic regularizer with hyperparameter *β* to update the VAE space dynamically such that it captures both cellular variations and the developmental dynamics of the scRNA-seq data. We test `scNODE` predictions when using different *β* ∈ *{*0.25, 0.5, …, 10.0*}*. Table S6 shows that the value of *β* affects model prediction accuracy, but `scNODE` is robust relative to the value of *β*. Moreover, `scNODE` performs better than baseline models in most cases. In experiments for the main text, we tune *β* values for `scNODE` in each case.

**Figure S7:**
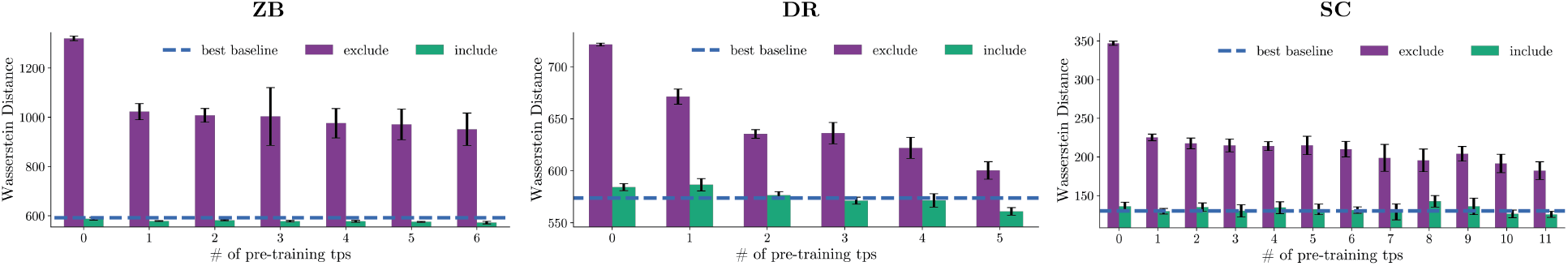
Compare Wasserstein distances of `scNODE` predictions in hard tasks when including and excluding the update of latent space. Dotted lines indicate the best baseline performance.

**Table S6:** Wasserstein distance of `scNODE` predictions when varying the regularization coefficient *β*. We report Wasserstein distance averaged over testing timepoints.

| Dataset/Task | Regularization Coefficient ($\beta$) | | | | | | | | Best Baseline |
| --- | --- | --- | --- | --- | --- | --- | --- | --- | --- |
|  | 0.25 | 0.5 | 0.75 | 1.0 | 2.5 | 5.0 | 7.5 | 10.0 |  |
| **ZB/easy** | 428.54 | 432.30 | 432.64 | 430.98 | 431.24 | 433.35 | 436.32 | 437.24 | 450.91 |
| **ZB/medium** | 649.85 | 651.64 | 647.88 | 644.91 | 646.56 | 645.86 | 635.19 | 636.29 | 666.45 |
| **ZB/hard** | 571.38 | 572.96 | 570.01 | 569.44 | 567.20 | 571.13 | 569.19 | 569.03 | 590.89 |
| **DR/easy** | 413.71 | 416.57 | 415.09 | 415.85 | 416.21 | 418.18 | 420.73 | 421.04 | 416.76 |
| **DR/medium** | 637.07 | 630.52 | 604.50 | 629.35 | 594.94 | 606.72 | 597.09 | 590.90 | 621.15 |
| **DR/hard** | 559.99 | 557.36 | 559.44 | 564.38 | 561.52 | 558.55 | 563.30 | 563.52 | 574.02 |
| **SC/easy** | 102.66 | 103.95 | 105.62 | 106.58 | 106.47 | 108.04 | 106.92 | 108.40 | 157.6 |
| **SC/medium** | 125.81 | 128.40 | 127.15 | 122.61 | 126.36 | 126.36 | 129.69 | 128.83 | 160.58 |
| **SC/hard** | 113.02 | 114.71 | 113.84 | 126.61 | 121.65 | 117.77 | 118.91 | 120.66 | 130.42 |

### S6.4 scNODE predictions help recover cell trajectories

We validate that `scNODE` can aid cell trajectory inference using the hard task of all three datasets. Specifically, we leave out several timepoints (see hard tasks in Table S1) and let `scNODE` and baseline model predict them back, in order to see whether model predictions can help with reconstructing cell developmental trajectories. We apply partition-based graph abstraction (PAGA) (Wolf et al., 2019) to construct cell trajectories. PAGA is a method that computes the topological structure of cell populations and represents population structures in interpretable graphs. Specifically, we use Louvain clustering (Blondel et al., 2008) to first cluster cells in the 2D UMAP space and apply PAGA to construct a structural graph of these clusters. We use *Scanpy* for Louvain clustering.

In addition, we use the Ipsen-Mikhailov (IM) distance (Ipsen, 2004) to quantitatively measure the similarity between the cell trajectory graphs constructed in each case. IM(*𝒢*_1_, *𝒢*_2_) is a graph similarity measurement defined as the square-root difference between the Laplacian spectrum of graphs *𝒢*_1_ and *𝒢*_2_. It ranges from 0 to 1, where 0 indicates maximum similarity between two graph structures and 1 indicates maximum dissimilarity. We use *nedtrd* package (McCabe et al., 2020) to compute the IM distance. Fig. S8 shows that `scNODE` predictions help recovering cell trajectories as IM(*𝒢*_true_, *𝒢*_scNODE_) *<* IM(*𝒢*_true_, *𝒢*_removal_). Moreover, IM(*𝒢*_true_, *𝒢*_scNODE_) being smaller than IM(*𝒢*_true_, *𝒢*_MIOFlow_) and IM(*𝒢*_true_, *𝒢*_PRESCIENT_) in all cases implies that `scNODE` predictions for missing timepoints best help to infer cell trajectories.

### S6.5 scNODE in perturbation analysis

We use ZB data to test `scNODE` ‘s ability of helping *in silico* perturbation analysis. We train `scNODE` with all timepoints of ZB data and construct the cellular state path in the latent space. The path is constructed with least action path (LAP), which has been used in previous works (Qiu et al., 2022, 2012; Wang et al., 2014) to construct cell fate transitions. The LAP method aims at finding the optimal path between two cell states while minimizing its action and transition time. Specifically, given starting point **X**_0_ and end point **X**_*K*_, LAP fins a path discretized as a sequence of *K* points *𝒫* = {**X**_0_, …, **X**_*K*_}. For each segment constrained between **X**_*k*−1_ and **X**_*k*_, its tangential velocity is defined as 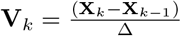 where Δ is the timestep taken by cells from **X**_*k*−1_. Therefore, we define the action *𝒮* along the path *𝒫* as

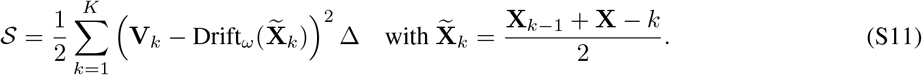

**Figure S8:**
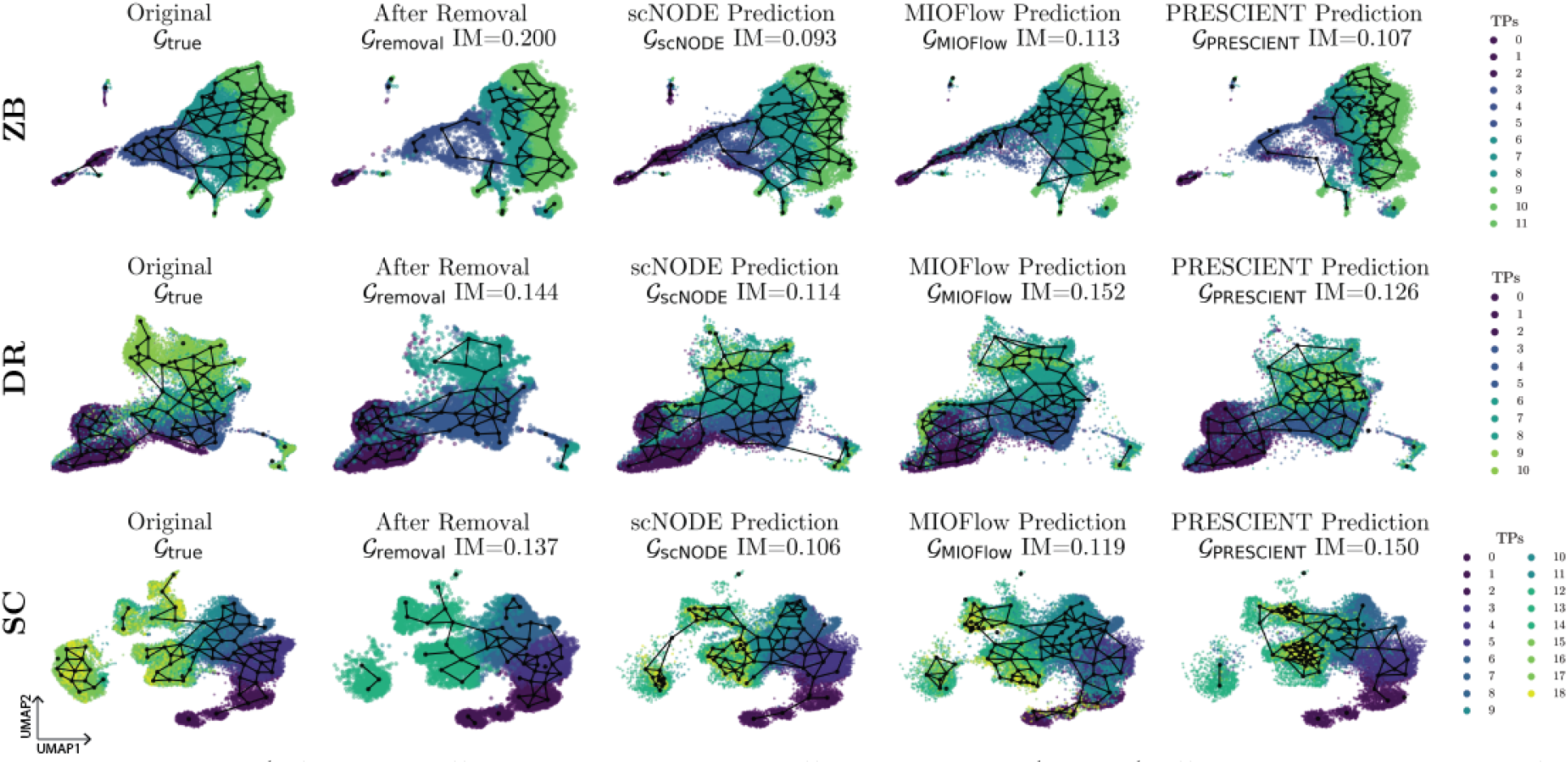
`scNODE` helps infer cell trajectories. Here are cell trajectories of data with all timepoints, after removal of timepoints in the hard task, and with model predictions. The connective structure is constructed with PAGA, where black nodes represent cell clusters and edges connect two nodes if their expressions are similar. We show the IM index between *𝒢*_true_ and the corresponding graph.

Here, LAP method aims to align the tangential velocity **V**_*k*_ with the differential velocity 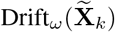 learned by `scNODE`, while having the least transition time. Therefore, the optimal path is

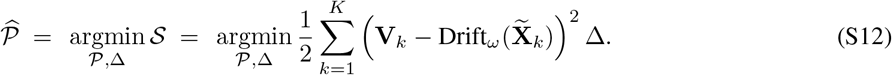

Solving Eq. S12 consists of two iterative steps

(1) Minimize action by fixing path *𝒫* and varying the timestep Δ through

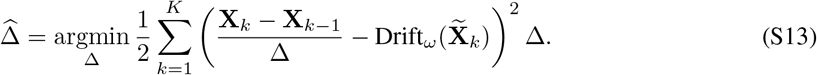

(2) Minimize action by fixing timestep 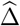 and varying path *𝒫*

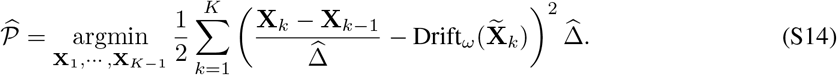

The starting (**X**_0_) and end point (**X**_*K*_) are fixed in the optimization.

We use *scipy*.*optimize*.*minimize* to solve these two objective functions. In our experiments, we construct two paths between the first timepoint and two cell populations (PSM and Hindbrain) with *K* = 8 (Fig. S9A). We set the starting point as the center of cells at the first timepoint (t=0) and the endpoint as the center of the cell population. We initialize timestep Δ = 1 and *𝒫* as equally spaced points from the starting to end points.

When finding the differentially expressed (DE) genes, we augment the LAP path with its nearest neighbors in the latent space (Fig. S9). Specifically, assuming ***𝒫*** = *{***X**_0_, …, **X**_*K*_*}* is the LAP path from **X**_0_ to **X**_*K*_, we have only 8 cells on the path which is insufficient for DE detection. Therefore, for each **X**_*k*_ ∈ *𝒫*, we find its nearest neighbors in the latent space in order to augment the path. We use *sklearn*.*neighbors*.*NearestNeighbors* to search for 10 nearest neighbors. The number of nearest neighbors affects DE detection, where fewer neighbors may be insufficient for good DE detection, and more neighbors may introduce noise. We select 10 neighbors for a proper tradeoff between efficiency and accuracy. Then we can use *Scanpy* to detect DE genes for the augmented path with the Wilcoxon rank-sum test.

Lastly, we perturb the expression profile of key genes in all cells at the starting timepoint by multiplying their expression values with different levels of coefficient *{*10^−3^, …, 10^3^*}* to mimic overexpressing and knocking-out, and let `scNODE` predict trajectories for the perturbed gene expression. The perturbations are expected to result in changes in cell fates. Specifically, we overexpress TBX16 (DE gene of PSM) or SOX3 (DE gene of Hind-brain). We classify predicted cells with a Random Forest (RF) classifier trained with all unperturbed cells. We use *sklearn*.*ensemble*.*RandomForestClassifier* with default parameters as the classifier and train it with all unperturbed cell expression as well as cell type labels in original datasets. We compare cell ratios when perturbing DE gene and random non-DE gene and find overexpressing TBX16 expression results in the increment of PSM cell ratios (Fig. S9**B**) to over 70%. Moreover, the ratio of Hindbrain cells decreased when overexpress SOX3 expressions (Fig. S9**C**). These indicate that detected DE genes are related to developmental dynamics. In applications of `scNODE`, these perturbations can involve multiple genes, use different cell populations, or be carried out at different timepoints.

**Figure S9:**
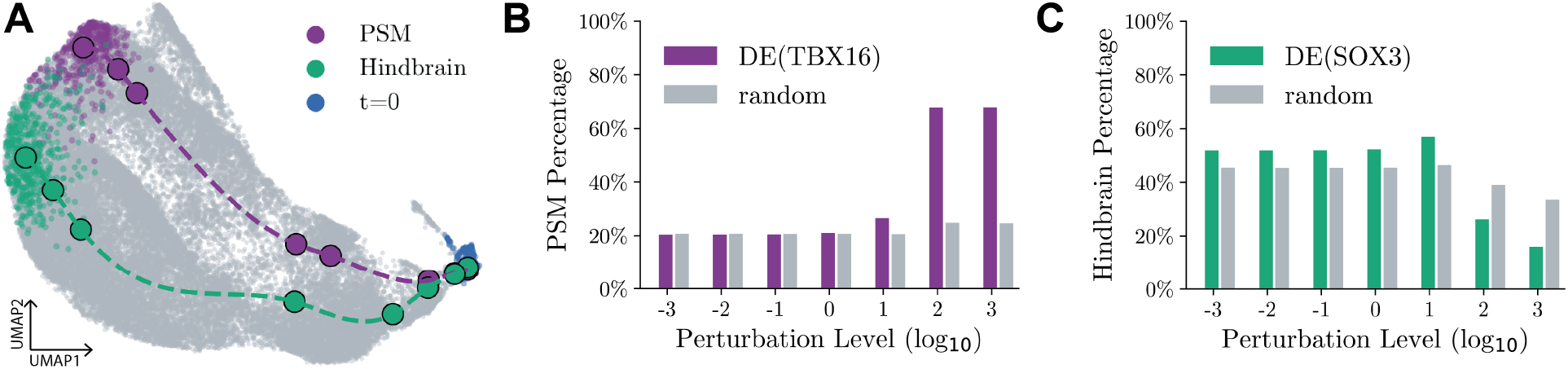
`scNODE` perturbation analysis results. (**A**) 2D UMAP visualization of the least action path between cells at the starting point (*t* = 0) and PSM/Hindbrain cell populations. (**B**) The ratio of PSM cells in predictions of different perturbation levels when perturbing TBX16 or ten random non-DE genes. (**C**) The ratio of Hindbrain cells in predictions of different perturbation levels when perturbing SOX3 or ten random non-DE genes.

